# Human condensin I and II drive extensive ATP–dependent compaction of nucleosome–bound DNA

**DOI:** 10.1101/683540

**Authors:** Muwen Kong, Erin Cutts, Dongqing Pan, Fabienne Beuron, Thangavelu Kaliyappan, Chaoyou Xue, Ed Morris, Andrea Musacchio, Alessandro Vannini, Eric C. Greene

## Abstract

Structural maintenance of chromosomes (SMC) complexes are essential for genome organization from bacteria to humans, but their mechanisms of action remain poorly understood. Here, we characterize human SMC complexes condensin I and II and unveil the architecture of the human condensin II complex, revealing two putative DNA–binding sites. Using single–molecule imaging, we demonstrate that both condensin I and II exhibit ATP-dependent motor activity and promote extensive and reversible compaction of double–stranded DNA. Nucleosomes are incorporated into DNA loops during compaction without being displaced from the DNA, indicating that condensin complexes can readily act upon nucleosome fibers. These observations shed light on critical processes involved in genome organization in human cells.

**One Sentence Summary:** ATP–dependent DNA compaction by human condensin complexes.

## Main Text

SMC (structural maintenance of chromosomes) complexes are essential for higher–order chromosome organization in all three domains of life (*1–4*). Mitotic chromosomes can be reconstituted through the actions of SMC condensin complexes (*5, 6*), thus defining condensin as a principle driver of chromosome architecture (*7, 8*). Higher eukaryotes have two conserved condensin complexes, condensin I (CI) and condesin II (CII), which share SMC subunits (SMC2 and SMC4) (*9*). The SMC subunits contain two ATP–binding head domains located at distal ends of two long (~50–nm) coiled–coil arms, the opposite ends of which dimerize to form the hinge domain (*10*). CI and CII have distinct non–SMC regulatory subunits, including a kleisin subunit (CAP–H and CAP–H2, respectively), which bridge the SMC head domains (*11*), and a pair of HEAT repeat subunits (CAP–D2/G and CAP–D3/G2, respectively), which can interact with DNA (*12, 13*) (Fig. 1A).

**Fig. 1.**
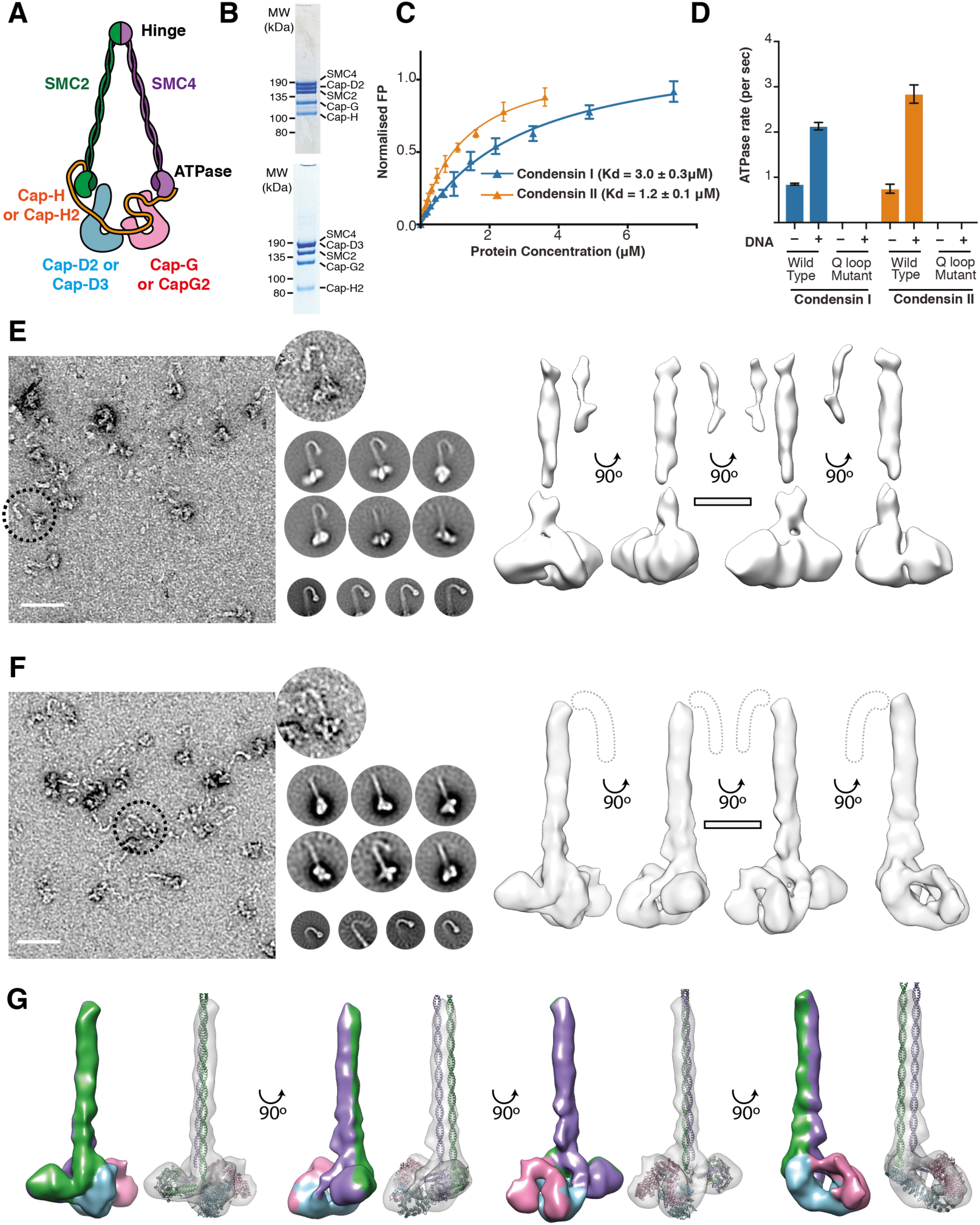
Biochemical characterization of human condensin I and II. (A) Schematic representation of human condensin complexes. (B) Analysis of purified CI and CII; Coomassie stained SDS PAGE gel from major peak from gel filtration with a Superose 6 10/300 column (MW, molecular weight). (C) Quantification of CI and CII DNA binding affinity using fluorescence polarization (FP) with 30 bp dsDNA labeled with 6FAM. (D) ATP hydrolysis activity of human CI and CII in the presence and absence of 100-fold excess of 50 bp of dsDNA, Q-loop mutants harbor ATPase site mutations SMC2 Q147L and SMC4 Q229L. (E) *left:* Negative stain electron micrograph of gradient crosslinked CI, the arrow indicates an example particle (scale bar in 50 nm). *Top Middle:* example 2D classifications of whole CI particles. *Bottom Middle:* 2D classifications, re-centered on the kink illustrating that while this feature is lost in some whole holocondensin classes, it is present in the raw data. *Right:* CI 3D model at 28.8 Å (scale bar 10 nm). (F) as in (E) but for CII, *Right:* CII 3D model at 20.4 Å, dashed line indicates hinge density not present in 3D model. (G) Pseudo-atomistic model of CII, with subunits and surface colored as in (A).

Two non-mutually exclusive models describe how condensins might compact DNA, the loop capture model, where condensin passively bridges distal chromosomal sites; and the loop extrusion model, where condensin actively extrudes loops of DNA Single molecule studies have shown that *Saccharomyces cerevisiae* condensin is an ATP-dependent motor protein that can extrude loops of DNA, providing support for the loop extrusion model (*15, 16*). Whether condensin from other organisms also act as DNA motors is unclear. Furtheremore, it is also unclear how condensin motor activity might is affected by nucleosomes, the basic structural unit of eukaryotic genomes. To address these fundamental questions, we sought to establish assays for observing the behaviors of the two distinct human condensin complexes on nucleosome–bound DNA molecules in real–time at the single molecule level.

Human CI and CII holocomplexes, each composed of five subunits, were recombinantly produced in insect cells. The complexes eluted as single peaks in gel filtration, and sample purity was confirmed by SDS page and mass-spectroscopy (Fig. 1B, Fig. S1A, Table S1). Both human CI and CII bind DNA and hydrolyze ATP. Fluorescence anisotropy assays (Fig. 1C) and electrophoretic mobility shift assays (EMSAs, Fig. S1B) indicate that CI has at least a 2-fold lower affinity for DNA (*K_d_* = 3.0 ± 0.3 µM) compared to condensin II (*K_d_* = 1.2 ± 0.1 µM). CI and II hydrolyzed ATP at rates of 0.85 ± 0.02 and 0.75 ± 0.10 ATP molecules per second per holocomplex, respectively. As expected, mutations of the Q-loop in the ATPase active sites (SMC2 Q147L and SMC4 Q229L,(*17*)) abolished ATPase activity. ATP hydrolysis was stimulated 2.5-and 3.8-fold by dsDNA for CI and II, respectively (Fig. 1D). Furthermore, both CI and II bind nucleosomes prepared with a 183-bp DNA substrate with an affinity similar to naked DNA but binding is severely impaired with a 147-bp DNA substrate (Fig. S1C), indicating that condensin complexes do not bind to core nucleosomes but prefer the flanking DNA.

Human condensin holocomplexes were analyzed by negative stain electron microscopy. To trap a specific conformational state, CI and II were pre-incubated with ATPγS, then separated and crosslinked using the GraFix method (*18*). CI and II samples appeared as ~60 nm elongated particles (Fig. 1E and F). The hinge, coiled-coil and HEAT domains (*7*) were clearly recognizable in micrographs and 2D classifications. Focused 2D classification was performed on the hinge, illustrating a rod-shaped fold, with a toroidal hinge domain and a kink is present ~15 nm from the hinge, consistent with previous studies on the SMC hinge (*19–22*). 3D EM maps of the human CI and II holocomplexes were obtained at ~28.8 and 20.4 Å, respectively. The resolution and the features of the map of CII were sufficient to fit homology models of SMC2/4 and CAP-D3/G2 (Fig. 1G). To independently validate domain contacts, crosslinking mass-spectroscopy was performed in the presence of ATPγS (Fig. S2D). The CII model satisfied 91.4% of crosslinked pairs present in the model, with the unsatisfied crosslinks suggesting an alternate conformation where the HEAT repeat domains contact the ATPase heads. The CII holocomplex displays two distinct compartments, lined with positively charged residues, which could each accommodate double stranded DNA (Fig. S3A-D). One compartment is formed by the engaged ATPase heads, resembling that observed in the ATPase domain of Rad50 (*23*) (Fig. S3E). The other compartment is formed by the kleisin and HEAT repeat domains, in agreement with what is described in *S. cerevisiae* cohesin (*24*) and *Bacillus subtilis* SmcScpAB (*25*).

To visualize DNA compaction by human condensin holocomplexes, we employed the single-molecule DNA curtain assay (Fig. 2A) (*26*). Addition of unlabeled condensin I (CI, 15 nM) or condensin II (CII, 20 nM) plus 4 mM ATP resulted in robust and progressive shortening of >35% of all DNA molecules (CI: N=110/276, CII: N=43/117). DNA compaction manifested in a localized increase in YOYO1 signal intensity that traveled towards the DNA tether point (Fig. 2B, Movie S1 and S2). The localized intensity typically translocated as a single bright spot, accompanied by the occasional appearance of weaker, elongated signal suggesting that a portion of the compacted DNA loop had been released (Fig. S4). Almost all compaction events were initiated at the free ends of the extended DNA molecules (CI, N=335/345, CII, N=215/215, Fig. S4), likely due to lower tension of the free DNA ends under flow (*27*), which would allow for transient loop capture as a prerequisite for productive compaction. Importantly, DNA compaction was strictly dependent upon ATP hydrolysis, as revealed in assays with either ATPase deficient condensin Q-loop mutants or nonhydrolyzable ATPγS (Fig. S5).

**Fig. 2.**
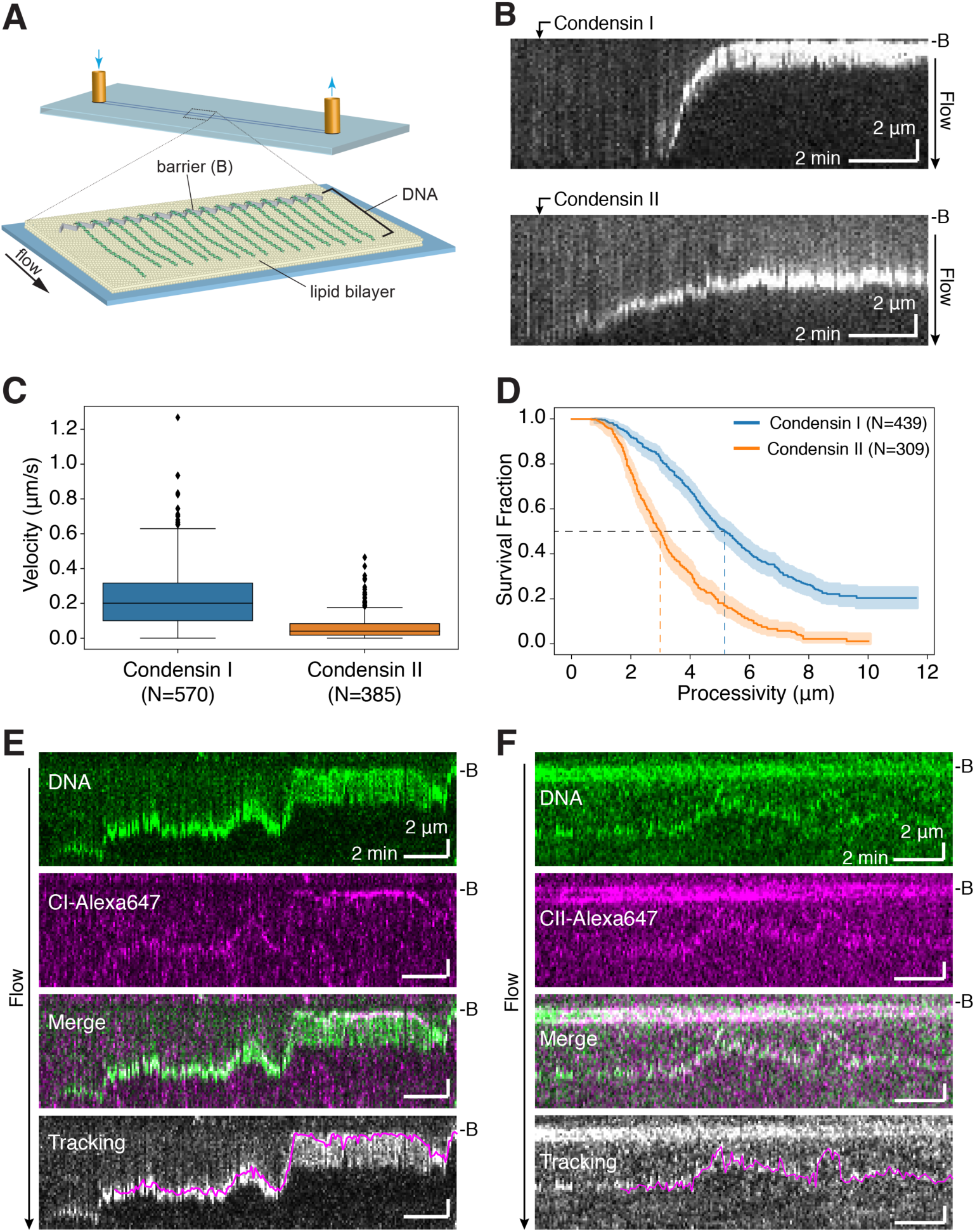
Real–time observation of DNA compaction by human condensin I and II. (A) Schematic of the single–tethered DNA curtain assay. (B) Kymographs showing compaction of single YOYO1–stained dsDNA molecules by human condensin I (*top*) and II (*bottom*). (C) Box plot of compaction velocities of CI and CII on naked DNA. (D) Kaplan-Meier estimated survival functions of compaction processivities on naked DNA. Shaded areas indicate 95% confidence intervals. (E) and (F) Kymographs showing compaction of single YOYO1–stained dsDNA molecules by Alexa647-labeled human CI and CII, respectively. Bottom panels show overlays of tracked trajectories of labeled condensins (magenta) with DNA signal.

Quantification of the DNA compactions yielded velocities 0.201 (0.216) µm/s (median and interquartile range, IQR), same below; or ~908 bp/s) and 0.040 (0.066) µm/s; (or ~182 bp/s) for CI (N=570) and CII (N=385), respectively (Fig. 2C). CI and CII processively compacted ~5.2 µm (~23.5 kbp; N=439) and 3.0 µm (~13.6 kbp; N=309) of DNA, respectively (Fig. 2D). The median velocities of human condensins were 2- to 10-fold higher than yeast and *X. laevis* condensins (*15, 17, 28*). The initiation of compaction was highly stochastic (Fig. S6) and lag times prior to compaction were distributed over the entire ~20 min observation period, consistent with the low *K_d_* values measured in bulk biochemical assays (Fig. S7A and S7B). Moreover, DNA molecules undergoing active compaction were often suddenly released, likely due to the spontaneous dissociation of condensin, allowing the DNA to return to its fully extended length (Fig. S4),. These observations demonstrate that the condensin-mediated compaction events were readily and completely reversible.

Our data suggested that condensin should co-localizes with the compacting loops of DNA. To test this prediction, we enzymatically conjugated Alexa647 fluorophore to both purified and ybbR-tagged CI and CII homocomplexes (Fig. S8). Fluorescently labeled CI and CII were observed to co-localize and track with the YOYO1 signal puncta formed during compaction indicating that CI and CII were directly localized with the compacting loops of DNA (Fig. 2E and 2F, Movie S3 and S4). Moreover, the Alexa647 signal did not increase appreciably during DNA compaction, indicating that compaction itself was driven by a finite number of condensin complexes that bound stochastically to the DNA (Fig. 2E and 2F). Compaction velocities and processivities of labeled condensins were found comparable to that of unlabeled proteins, while photobleaching experiments revealed that the majority (>70%) of compaction events were driven by a single labeled CI complex (Fig. S9). Furthermore, using either labeled or unlabeled condensins on double-tethered DNA curtains, we observed ATP-dependent linear translocation of human condensins, similar to what we have reported for yeast condensin (Fig. S10), indicating that the intrinsic motor activity of condensin is conserved from yeast to human (*15*).

Active loop extrusion has emerged as a widely accepted model for DNA compaction by condensins (*14, 29–32*). We sought to visualize the looping process for unlabeled human condensins using YOYO1-stained DNA molecules that were tethered in a U-shaped configuration (Fig. 3A). For both condensin I and II, DNA puncta were observed first forming at the distal end of the U-shaped DNA and then moved progressively towards the DNA tether points and this compaction activity was strictly dependent upon ATP-hydrolysis (Fig. S11). These observations indicate that a DNA loop is formed as human condensins translocate on DNA, as previously reported for yeast condensin (*16*).

**Fig. 3.**
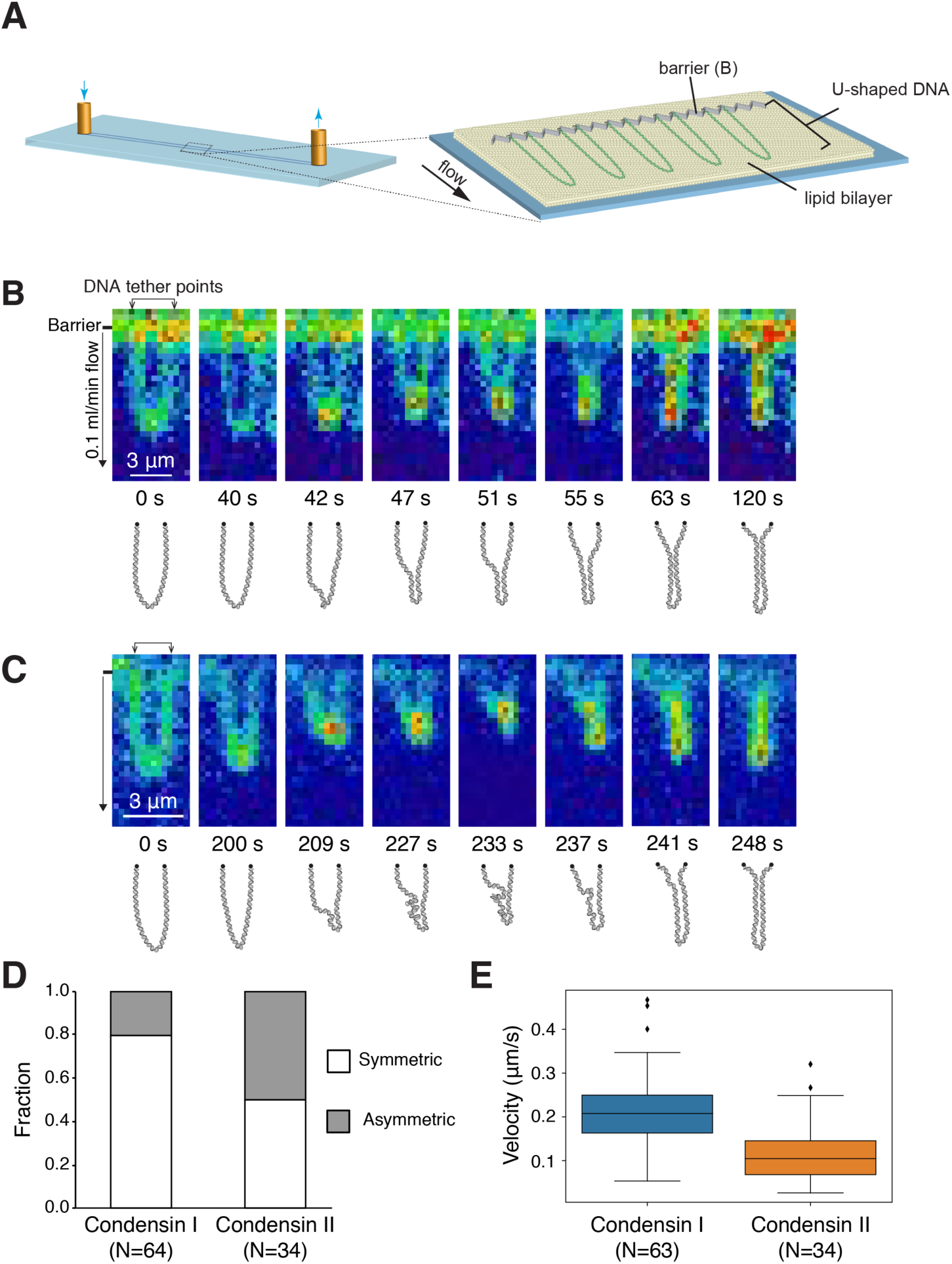
Loop extrusion of U–shaped DNA molecules by human condensin I and II. (A) Schematic of U–shaped DNA curtain assay. (B) Snapshots and schematics of loop extrusion of YOYO1–stained U–shaped DNA molecules by human CI (*top*) and CII (*bottom*). (C) Bar graph comparing the number of symmetric and asymmetric compaction events. (D) Box plot of loop extrusion velocities.

The manner in which the DNA was compacted could be categorized as either symmetric, where condensin appeared to act on both sides of the U-shaped DNA molecules (Fig. 3B, *top*; Fig. S12A, and Movie S5), or asymmetric, where condensin appeared to act on just one side of the DNA (Fig. 3B, *bottom*; Fig. S12B, and Movie S6). We classified events according to the symmetry in which the DNA puncta progressed and found that 80% of all events involving CI were symmetric with 20% being asymmetric, while equal numbers of symmetric and asymmetric events were observed with CII (Fig. 3C). These observations contrast with yeast condensin, which exhibit 100% asymmetry in loop extrusion (*16*). We quantified the progression of DNA puncta as a measure of loop extrusion rates, obtaining median velocity values comparable to those of single-tethered DNA compaction events (CI: 0.208 (0.087) µm/s or ~1056 bp/s; CII: 0.105 (0.077) µm/s or ~535 bp/s, Fig. 3D). These rates are consistent with those previously reported for yeast condensin loop extrusion, which varied between 200 bp/s to 1000 bp/s (*16*).

Nucleosomes serve as the first layer of compaction in the 3D organization of eukaryotic genomes (*33*). Therefore, we investigated whether nucleosomes may influence the compaction process by human condensins. To this end, we assembled ATTO-647N-labeled histone octamers on λ-DNA under conditions that yielded ~3-4 nucleosomes per DNA (Fig. S13). On single-tethered DNA curtains (Fig. 4A), we observed extensive compaction of nucleosome-bound DNA molecules by both condensin I and II (Fig. 4B, Movie S7 and S8). Nucleosomes were readily incorporated into the compacted DNA during this process, as was evident by the co-localization of ATTO-647 and YOYO1 signal as the punctum traveled towards the barrier (Fig. 4B). Upon encountering individual nucleosomes, approximately 80% of all compaction events proceeded without stopping, releasing, or reversing direction (CI: N=336/397, CII: N=141/178, Fig. S14). Most (~90%) of these encounters did not exhibit any pausing within our temporal resolution (5 sec, CI: N=304/336, CII: N=122/141), indicating that the nucleosomes did not hinder either CI or CII movement. Furthermore, neither the velocity nor processivity of compaction was significantly affected by the presence of nucleosomes for either condensin I or II (Fig. S15). More importantly, the inherent reversibility seen in these compaction reactions unequivocally demonstrated that compaction did not displace the nucleosomes from the DNA or alter their positions on DNA within our resolution (Fig 4B).

**Fig. 4.**
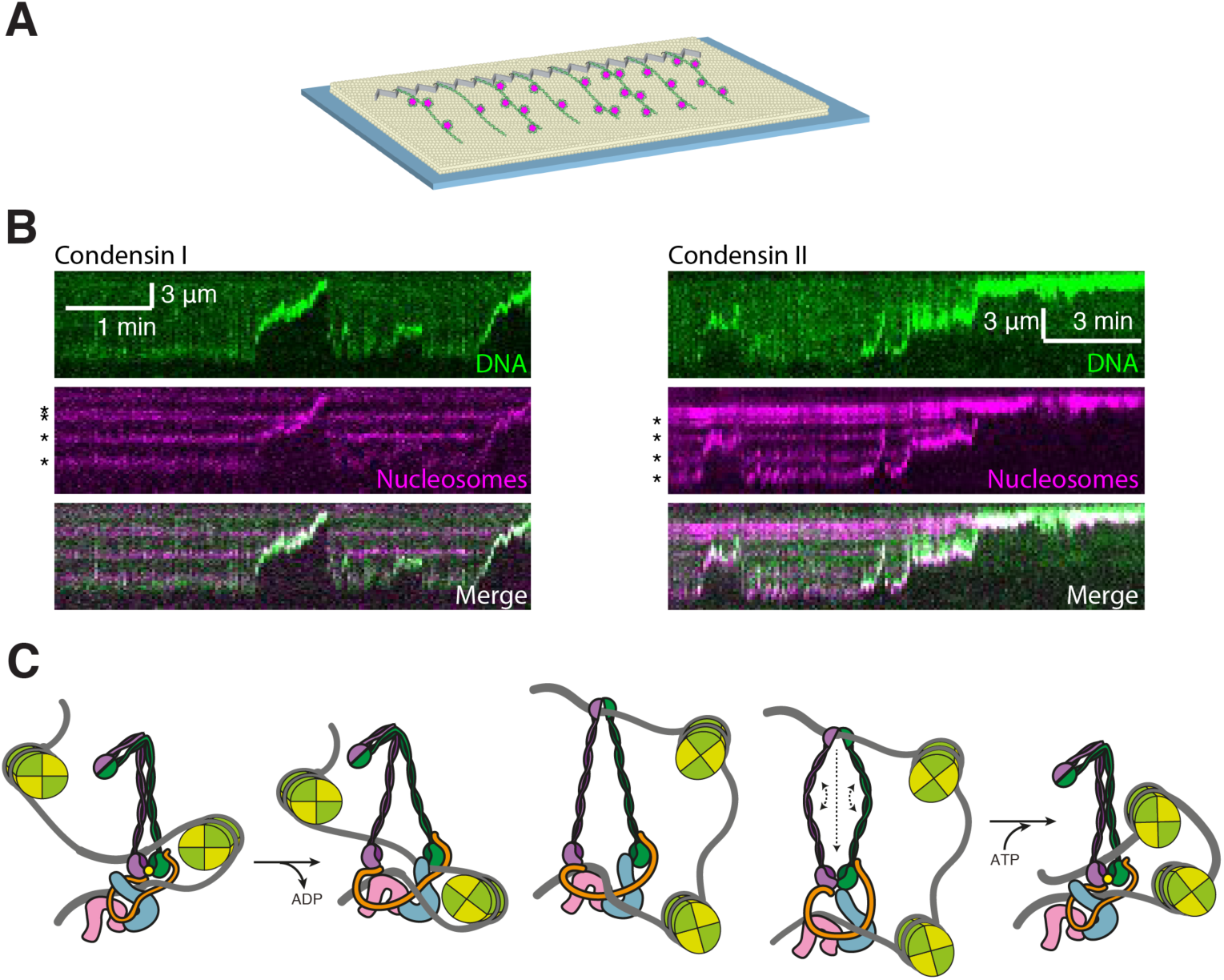
Compaction of nucleosome–bound DNA substrates by human condensin I and II. (A) Schematic of single–tethered DNA curtain assays with nucleosome (magenta)–bound DNA substrates. (B) Kymographs showing CI (*left*)– and CII (*right*)–mediated compaction of single–tethered YOYO1–stained DNA (green) bound by ATTO647–labeled nucleosomes (magenta). (C) Model for compaction of nucleosome–bound DNA by human condensins. Dashed lines indicate possible movements.

Our data reveals the general architecture of a condensin holocomplex and shows that the ATP-dependent mechanism of DNA compaction is broadly conserved among eukaryotes. Furthermore, the single molecule experiments support a model in which human condensins readily bypass nucleosomes and incorporate them into DNA loops, suggesting that condensin complexes could act through similar ATP-dependent motor-based mechanisms while compacting chromosomes in crowded physiological settings. Our EM model indicates there are channels that could accommodate dsDNA, however these are not large enough to allow passage of a nucleosome. Based on our protein crosslinking data and previous observation that the SMC2/4 arms can open (*34*), we speculate that condensin can step over nucleosomes during DNA compaction (Fig. 4C). The coiled-coil arms of condensin SMC2/4 measure ~50 nm in length and could easily allow the ~10 nm chromatin fiber to pass through the ring of a topologically bound condensin holocomplex. Although many details remain to be elucidated, several models currently exist to describe the underlying mechanism of condensin-mediated loop extrusion. Inchworm-like models that involve either scrunching of the coiled-coils concurrently (*14*) or the alternating opening and closing of the two (*32, 35*) are consistent with our observations and can allow for large step sizes.

## Acknowledgments

We thank members of the Greene, Vannini and Musacchio groups for discussion throughout this study. We thank Jyoti Choudhari and the mass spectrometry groups at the Institute of Cancer research for measuring confirming condensin sample composition and at the Max Planck Institute of Molecular Physiology for measurement and data analysis of cross-linked condensin complexes.

## Funding

This work was funded by a MIRA grant from the National Institutes of Health to E.C.G. (R35GM118026). A.V. is supported by a Cancer Research UK Programme Foundation (CR-UK C47547/A21536) and a Wellcome Trust Investigator Award (200818/Z/16/Z). A.M. gratefully acknowledges funding by the Max Planck Society, the European Research Council (ERC) Advanced Investigator Grant RECEPIANCE (proposal 669686)

## Author contributions

M.K. performed DNA curtain experiments and data analysis. E.C. purified human condensin complexes, performed and analyzed biochemical assays and structural characterization and modelling. D.P. performed crosslinking mass-spectroscopy experiments. F.B. assisted in electron microscope sample preparation, data collection and data processing. E.M. assisted with electron microscope data processing. T.K. made nucleosomes for EMSAs. C.X. prepared histones used in DNA curtain experiments. M.K., E.C., A.V. and E.C.G. designed and supervised research and co-wrote manuscript.

## Competing interests

Authors declare no competing interests.

## Data and materials availability

All data are in the paper and supplementary materials will be made available upon request.

## Supplementary Materials

### Materials and Methods

#### Expression, purification, and labeling of human condensin complexes

All constructs purified and used are listed (Table S1). The subunits of human condensin I and II, sub–complexes and Q–loop mutations and were assembled into biGBac vectors (*1*). Viral bacmids were generated from biGBac vectors using Tn7 transposition in DH10EMBacY cells, transfected into Sf9 cells using cellfectin II (Invitrogen) and resultant virus harvested after 3 days. Virus was further amplified in Sf9 cells and protein expressed in HighFive cells, which were harvested by centrifugation 3 days after infection. Cell pellets were resuspended in condensin purification buffer (20 mM HEPES [pH 8], 300 mM KCl, 5 mM MgCl_2_, 1 mM DTT, 10% glycerol) supplemented with 1 Pierce protease inhibitor EDTA–free tablet (Thermo Scientific) per 50mL and 25 U/mL of Benzonase (Sigma) and lysed with a dounce homogeniser followed by brief sonication. Lysate was cleared with centrifugation, loaded on to a StrepTrap HP (GE), washed with condensin purification buffer and eluted with condensin purification buffer supplemented with 5 mM Desthiobiotin (Sigma). Protein containing fractions were pooled, diluted 2-fold with Buffer A (20 mM HEPES [pH 8], 5 mM MgCl_2_, 5% glycerol, 1 mM DTT), loaded on to HiTrap Heparin HP column (GE), washed with Buffer A with 250 mM NaCl, then eluted with buffer A with 500 mM NaCl. Finally, size exclusion chromatography was performed using Condensin purification buffer and a Superose 6 16/70 or increase 10/300 column (GE) (Fig. S1A). Purified condensin I and II complexes were analyzed by SDS page (Fig. 1B) and MS/MS, indicating they were the major species in each sample and all subunits were present (Table S2).

Labelled condensin I and II; wild type and corresponding Q–loop mutants were labelled using the SFP transferase to couple a CoA conjugated Alexa647 (Thermo fisher) via a ybbR tag at the C–terminus of CAP-H in the case of Condensin I, or in the N–terminus of SMC4 in the case of Condensin II. Purification of SFP and labelling was performed as in (*2*), except that the conjugation CoA with Alexa647–maleimide was performed in 5–fold excess of Alexa647–maleimide, quenched with DTT in 10–fold excess over Alexa647–maleimide and used directly in reactions. Excess unconjugated material was separated from condensin complexes using size exclusion chromatography with a Superose 6 10/300 column and the presence of all subunits and specific conjugation confirmed by SDS page (Fig. S8A–D). EMSAs were performed on labelled material to confirm the fluorophore did not affect DNA binding (Fig. S8E).

#### Nucleosomes for Gel Shift Analysis

The Widom 601 (*3*) DNA sequence (147 bp underlined) alone and with 36 bp of linker DNA (183 bp DNA), as follows, was used to make nucleosomes: 5’<u>ATC GAG AAT CCC GGT GCC GAG GCC GCT CAA TTG GTC GTA GAC AGC TCT AGC ACC GCT TAA ACG CAC GTA CGC GCT GTC CCC CGC GTT TTA ACC GCC AAG GGG ATT ACT CCC TAG TCT CCA GGC ACG TGT CAG ATA TAT ACA TCC GAT</u> TAA CGA TGC TGG GCA TAA GCG TGG TTC AAT ACC GGC 3’. DNA was generated using large–scale PCR with in house *Pfu* polymerase, and labeled primers (IDT). The obtained PCR products were pooled, and ethanol precipitated. The pellets were re–dissolved in buffer A (10 mM Tris–HCl [pH 8], 1 mM EDTA [pH 8]) and loaded into a Mono Q 5/50 GL ion–exchange column (GE Healthcare) and eluted with a salt gradient in buffer B (10 mM Tris–HCl [pH 8], 2 M NaCl, 1 mM EDTA [pH 8]). Fractions corresponding to the DNA fragments were assessed by 4–12% polyacrylamide gel, pooled, followed by ethanol precipitation. The pellet was re–dissolved in buffer A, and stored at –20°C.

Expression, purification and assembly of human histones H2A, H3, H4 and *Xenopus laevis* H2B was performed as described previously (*4*). Briefly, histone proteins were expressed in *E. coli*, and purified from inclusion bodies using cation and anion ion–exchange chromatography, then subsequently lyophilized for long–term storage at –20°C. Individual lyophilized histones were mixed at 1.2 fold excess of H2A and H2B and resuspended in unfolding buffer (20 mM Tris–HCl pH 7.5, 7 M guanidine hydrochloride, 5 mM DTT) for 45 minutes at room temperature, and dialyzed against refolding buffer (10 mM Tris [pH 7.5], 2 M NaCl, 1 mM EDTA, 5 mM β–mercaptoethanol) for 18 h at 4°C, and then subjected to size–exclusion chromatography on a S200 16/60 gel filtration column (GE Healthcare) equilibrated in refolding buffer. The fractions corresponding to histone octamers were pooled, concentrated and flash frozen in liquid nitrogen and stored at –80 °C. The nucleosomes were reconstituted by salt dialysis method as described previously (*5*). Purified DNA and histone octamer were mixed at 1:1.1 ratio of DNA:octamer in a high salt buffer (10 mM Tris-HCl pH 7.5, 2 M NaCl, 1 mM EDTA pH 8, 1 mM DTT) and the sample was dialyzed gradually to low salt buffer (10 mM Tris-HCl pH 7.5, 0.2 M NaCl, 1 mM EDTA pH 8, 1 mM DTT) over 24 h at 4 °C using a peristaltic pump. The sample was further dialyzed against in low salt buffer for 4 h, then in nucleosome buffer (20 mM Tris-HCl pH 7.5, 50 mM NaCl, 1 mM EDTA pH 8, 1 mM DTT) overnight at 4 °C, before being concentrated and stored at 4 °C

#### Gel shift analysis

Double stranded DNA for EMSAs were made by annealing single stranded DNA oligos using a temperature gradient from 95 to 4°C. DNA oligos were purchased with 5’ Cy5 fluorophore or 6–FAM on the reverse strand (IDT), with the 30 bp sequence composed of 5’–CTG TCA CAC CCT GTC ACA CCT GTC ACA C–3’, and the 36 bp sequence was 5’ TAA CGA TGC TGG GCA TAA GCG TGG TTC AAT ACC GGC–3’. EMSAs with unlabeled protein were performed by incubating 50 nM of Cy5 labelled DNA with indicated concentration of protein on ice for ~30 minutes in condensin purification buffer, before running 4 µL on 2% agarose gel in 0.5x Tris borate buffer (TB) for 30–60 minutes. EMSAs with Alexa647 labelled protein were performed by incubating 25 nM of 6FAM labelled 30 bp DNA with indicated concentration of protein on ice for ~30 minutes in condensin purification buffer, before running 6 µL on 2% agarose gel in 0.5x TB for 30 minutes. Gels were imaged using a Typhoon FLA 9000 scanner (GE) and analyzed with imageQuant. Nucleosome EMSAs were performed by incubating 50 nM of nucleosome or free dsDNA with indicated concentration of protein and running on 2% Agarose for 60 minutes.

#### ATP hydrolysis Assays

ATP reactions were performed at 37°C using 0.2 μM of protein in ATPase buffer (10 mM Tris [pH 7.5], 100 mM NaCl, 5 mM MgCl_2_, 0.5 mM DTT and 0.1 mg/ml BSA) in the presence or absence of 20 μM of 50 bp of annealed dsDNA. Reactions were started by adding 2 mM of cold ATP supplemented with 1 μCu of [α–^32^P] ATP (800Ci/mmol) (Perkin–Elmer) and aliquots were removed and quenched with 100 mM EDTA at multiple time intervals. Aliquots were spotted onto PEI cellulose F TLC plates (Merck) and run in 1 M formic acid and 300 mM LiCl. TLC plates were used to expose a phosphor–imager plate that was subsequently scanned on a Typhoon FLA 9000 scanner (GE) and analyzed with imageQuant. ATP hydrolysis rate was determined by linear fit of the ADP/ATP ratio during in the linear range of the reaction. ATP hydrolysis assay for each sample was performed with three replicates.

#### Fluorescence polarization

Fluorescence polarization experiments were performed by mixing 50 nM of 6–FAM labelled 30 bp dsDNA with indicated concentrations of protein in FP buffer (20 mM Tris [pH 7.5], 75 mM NaCl, 2.5% glycerol, 1 mM DTT), incubating at room temperature for 30 minutes before reading on a BMG labtech POLARstar Omega plate reader. Three replicates were performed for each protein concentration and globally fit using the following:

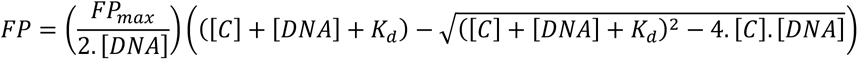

Where [C] and [DNA] are the concentration, in μM, of condensin and 6–FAM labelled DNA respectively, FP_max_ the maximum change in Fluorescence polarization in mAu and K_d_ is the equilibrium dissociation constant. Fit curves were plotted normalized by dividing by FP_max_, with error bars indicating standard deviation.

#### Negative Stain Electron microscopy

Samples for EM were prepared using gradient–fixation (Grafix) (*6*). Gradients were made by sequential freezing of Grafix buffer (50 mM HEPES [pH 8], 750 mM NaCl, 2 mM TCEP, 2 mM MgCl_2_) with 50%, 40%, 30%, 20% and 15% glycerol, and 0.2%, 0.1%, 0.1%, 0.05% and 0% glutaraldehyde respectively. The top three layers (15–30% glycerol, also contained ~40 µM of ATPγS). Condensin complexes pre–incubated for ~30 min with ~2 mM ATPγS were carefully applied to the top of thawed gradients and run at 29k rpm for up to 20 hours. Fractions containing crosslinked species were quenched with 100 mM Tris pH 8 and SEC run using Superose 6 10/300 column (GE) in EM buffer (20 mM HEPES [pH 8], 200 mM KCl). Negatively stained samples were prepared onto Quantifoil copper grids R1.2/1.3 supporting an additional layer of thin carbon prepared in house. The grids were glow discharged for 1 minute at 15 mA, the sample applied for 10 seconds and then washed twice with water before staining with 2% uranyl acetate for 1 minute. The grids were screened on a Tecnai T12 electron microscope operating at 120 kV. Micrographs were collected on a Tecnai TF20 (Thermo Fisher Scientific, USA) microscope operating at 200 kV using a 4k x 4k F416 CMOS detector (TVIPS Gmbh, Germany) with the EM–Tools automated data collection software (TVIPS Gmbh, Germany). Example micrographs of condensin I and II are shown (Fig. 1E and F). Data processing was done using RELION–3 (*7*).

The model for condensin II was determined first. Particles that were well stained, with obvious density for the coiled coil and heat domains were picked manually, resulting in a total of 3,637 particles. Poor particles were removed with subsequent rounds of 2D classification, resulting in 3,251 particles, which were used for 3D classification. A subset of 1,288 particles were used to generate an initial model. 3D classification was used to generate 3 classes, using the initial model as a reference. One class composed of 1565 particles yielded a model with well defined, domain–like features. After 3D refinement and post processing this model had a resolution of 20.4 Å. For condensin I, particle picking was performed as for condensin II, resulting in 6,601 particles. After multiple rounds of 2D classification, 5,977 particles were subjected to 3D classification, using the initial model used for condensin II. One class of 2,055 particles, that yielded a model with clear features, was further refined and post–processed to a final resolution of 28.8. The final particle set used for 3D classification was subjected to 2D classification (Fig. 1E and 1F) and 2D classes were compared to 3D models, resulting in reasonable agreement in features (Fig. S2A).

#### Chemical cross–linking and mass spectrometry (XL–MS)

Condensin I and II complexes were diluted in 200 µl buffer containing 20 mM HEPES [pH 8], 300 mM KCl, 10 mM MgCl_2_, 1% glycerol, 1 mM TCEP with addition of 2 mM ATPγS. Disuccinimidyl dibutyric urea (DSBU) was added to the solution to a final concentration of 3 mM and incubated at 25°C for 1 hour. The cross–linking reaction was terminated by adding Tris–HCl pH 8.0 to a final concentration of 100 mM and further incubation at 25°C for 30 min. Samples were taken before and after cross–linking reaction and examined by SDS–PAGE (Fig. S2C).

Protein digestion, peptide purification and MS analysis were performed as previously described (*8*). Cross–linked condensin complexes were precipitated by mixing with 800 µl acetone and incubation at –20°C for more than 16 hours. After removal of acetone, protein pellets were dissolved in 8 M urea containing 1 mM dithiothreitol (DTT). Chloroacetamide (5.5 mM) was used to alkylate the cysteine residues. Proteolysis was performed by Lys–C for 3 hours at 25°C and followed by trypsin for 16 hours at 25°C. Trifluoroacetic acid (TFA) was added to the solution to a concentration of 0.2% to stop the digestion. Peptides were purified by size–exclusion chromatography (SEC) using a Superdex Peptide 3.2/300 column (GE healthcare). Four SEC fractions (B8–B5) of each cross–linked sample were collected separately and measured twice by LC–MS/MS as previously described (*8*). Dried peptides were dissolved in water containing 0.1% TFA and were separated on the Ultimate 3000 RSLC nano system. Data were acquired using the Q–Exactive Plus mass spectrometer (Thermo Fisher Scientific) in data–dependent MS/MS mode. For full scan MS, we used mass range of m/z 300−1800, resolution of R = 140000 at m/z 200, one microscan using an automated gain control (AGC) target of 3e6 and a maximum injection time (IT) of 50 ms. Then, we acquired up to 10 HCD MS/MS scans of the most intense at least doubly charged ions (resolution 17500, AGC target 1e5, IT 100 ms, isolation window 4.0 m/z, normalized collision energy 25.0, intensity threshold 2e4, dynamic exclusion 20.0 s).

Program MeroX version 1.6.6.6 (*9*) was used for cross–link identification. In the settings of MeroX, the precursor precision and the fragment ion precision were changed to 10.0 and 20.0 ppm, respectively. RISE mode was used and the maximum missing ions was set to 1. A false discovery rate of 5% was used as the cut–off to exclude the candidates with lower MeroX scores. A mass–deviation range of −4 to 6 ppm was used to further exclude cross–link candidates with mass deviation outside of the range. Non–redundant cross–link lists (Table S3) only contain the identified cross–links with the highest MeroX scores if the same cross–links were identified more than twice. The total list of crosslinks for condensin I and II was filtered for those that are present at least 3 times in total and visualized using *xVis* (Fig. S2D) (*10*).

#### Structural Modelling

To generate a pseudo atomic model, the crystal structure of the human SMC2/4 hinge (PDB ID: 4U4P, SMC2 residues 507–673, SMC4 610–754) was combined with homology models of SMC2/4 ATPase domain (template: *S. cerevisiae* SMC3 4UX3, SMC2 residues 2–192 and 1006–1181 and SMC4 residues 83–278 and 1108–1284), CAP–D3 (template: human Pds5B 5HDT, residues 212–1261) and CAP–G2 (template: human CAP-G 5OQQ, 275–1111) generated using Phyre2 (*11*) and coiled coils built using CCBuilder2.0, using a radius of 5 nm, a pitch of 70 and an interface angle of 45 degrees (SMC2 residues 195–504 and 673–1003, and SMC4 residues 301–589 and 766–1087). ATPase heads were modelled together in the engaged conformation by aligning above mentioned homology models to ATPase domains in Rad50 crystal structure 5F3W (*12*).

Subunits were sequentially fit into condensin II map using *Chimera* (*13*), subtracting the fit density before the subsequent subunits were fit. The SMC2/4 ATPase heads were fit first, resulting in two likely fits in the density proximal to the coiled–coil region, as the SMC2/4 model had pseudo symmetry. As the N–terminal region of CAP-H2 is predicted to interact with SMC2 arm adjacent to the ATPase head (*14*), the fit which places the SMC2 coiled–coil arm in the region with most map density was selected. Next, the coiled–coil domain was fit, such that there would be sufficient space for un–modelled regions between the ATPase domains and the coiled–coils. This arrangement suggests there would be have to be considerable bend in the coiled–coils near the ATPase heads. Sequence alignment of human and *S. cerevisiae* SMC 1–4 and *E. coli* MukB sequences suggestion this region can contain a number of glycine and proline residues, which cause a break in helicity, demonstrated in the crystal structure of the *S. cerevisiae* SMC3 ATPase (PDB ID: 4UX3, Fig. S2B). Of the residual unaccounted map, one region had harp–shaped density, which was complementary for the shape of the CAP–G2 homology model. The remaining density was fit with the CAP-D3 model, which places the N–terminus of CAP–D3 near the coiled–coil region of SMC2 and the C–terminus near CAP–G2. The shape complementarity for CAP–D3 was poorer than for the rest of the structure, which is most likely due to a lack of directly homologous structures. The fit of CAP-D3 was improved by twisting residues 212–648 relative to the C–terminal region, such that it could follow the curve observed in the density.

While there was not sufficient structural information to model CAP–H2, the modelled subunit arrangement is consistent with interaction studies indicating the N–terminus of CAP–H2 binds to SMC2, then to CAP–D3, then CAP–G2 and has the C–terminus interacting with SMC4 (*14*).

#### Structural Analysis

The Condensin II model was then compared to mass–spec crosslinking data. Using *Xlink Analyzer* (*15*) with a distance cut–off threshold of 35 Å, 91.4% of crosslinked pairs present in the modelled structure were satisfied. The crosslinking data also supports the orientation of CAP–G2 and a considerable interaction between the N–terminal region of CAP–G2 and the C–terminal region of CAP–D3, as the crosslink from CAP–G2:340 to CAP–D3:1214 is the most abundant non–SMC inter–domain crosslinks. The engagement of the ATPase heads is also supported by crosslinks, such as those between SMC2:1079–SMC4:1267 and SMC2:12–SMC4:1187. There are multiple crosslinks between the SMC2/4 coiled–coils, and these are mostly enriched in region between the kink to the hinge, suggesting this region may close independently of the rest of the SMC arms. Examining the violated crosslinks, there is a cluster of violated crosslinks between the interface of CAP-G2/D3 with SMC2, and a crosslink between the hinge and coiled–coil region near the ATPase domain, which could represent an alternate conformation of the complex.

The condensin II map has two compartments, one formed by the engaged SMC heads, and another formed by the kleisin, HEAT repeat domains and the closed SMC arms (Fig. S3A). The vacuum electrostatics of the surface lining these compartments were calculated using PyMOL v1.8 (Schrödinger, LLC.), suggesting a predominantly positive surface that should allow for DNA binding (Fig. S3 B–D). Comparison between the condensin II model and the ATPase domain of Rad50 (5F3W (*12*) bound to DNA, suggests that DNA could be bound in this compartment via interaction with the SMC2/4 ATPase heads. Comparison of the *S. cerevisiae* CAP–G/H DNA bound structure, suggests that if this binding site is present in condensin II it would be outward facing (Fig. S3F).

The condensin I 3D reconstruction was not of sufficient resolution to allow unambiguous structural modelling; however the crosslinking data provides some structural insights. There are crosslinks made by CAP–H supporting previously proposed interactions between the CAP–H N–terminus and SMC2, CAP–H2 residues around 500 with CAP–G and the CAP–H C–terminus with SMC4 (*14, 16*). Crosslinks also show similarities with condensin II such as a similar enrichment of crosslinking between the coiled–coils of SMC2/4 from the kink to the hinge, interactions of the HEAT domains with the SMC ATPase domains and surrounding coiled–coil regions, and interactions between the hinge and HEAT domains, suggesting similar conformational changes occur in both condensin I and II.

#### Single–molecule experiments

Single–molecule DNA curtain assays were performed as previously described (*17, 18*). To prepare DNA substrates for curtain experiments, 1.6 pmol of λ–DNA (N3011S, New England Biolabs) was mixed with 100–fold molar excess of biotin– or digoxigenin end–modified oligonucleotides (IDT) in 1X T4 ligation buffer (B0202S, New England Biolabs) in a 600 μL reaction. Oligos BioL (5’Phos–AGG TCG CCG CCC–3Bio) and DigR (5’Phos–GGG CGG CGA CCT–3Dig_N) were used for substrate in single–tethered experiments, whereas BioL and BioR (5’Phos–GGGCGGCGACCT–3Bio) were used to generate the U–shaped DNA substrate. The mixture was assembled at 65 °C and incubated for 5 minutes before cooled down to room temperature. 5 μL of T4 DNA ligase (M0202S, New England Biolabs) was then added to the reaction and ligation was carried out overnight at room temperature. DNA substrate was precipitated by adding PEG8000 (Sigma, Cat. No. 89510) and MgCl_2_ to the final concentrations of 10% and 10 mM, respectively, and incubating at 4°C with rotation. Following centrifugation of sample at 14,000 x *g* for 5 minutes, resulting DNA pellet was washed once with 70% ethanol, resuspended in ddH_2_O, and stored at 4°C. Briefly, in a custom–made microfluidic flow cell, biotinylated DNA substrate was anchored to the supported lipid bilayer on the surface of a quartz slide with nanofabricated chromium barriers, through biotin–streptavidin linkage. Double–stranded DNA was stained with buffer containing 0.5 nM YOYO–1 (Y3601, Invitrogen) and visualized through 488 nm excitation (Coherent Sapphire) on a custom–built prism–type TIRF microscope (Nikon Eclipse TE2000–U) equipped with a 60x water immersion objective and an EMCCD camera (Photometrics Cascade II: 512 or Andor iXon X3). All experiments were carried out at 37°C in condensin buffer (40 mM Tris–HCl [pH 7.5], 125 mM NaCl, 5 mM MgCl_2_, 1 mM DTT, 0.5 mg/mL BSA), supplemented with either 4 mM ATP or ATPγS. Unlabeled condensin in condensin buffer was continuously injected into the flow cell via a syringe pump (KD Scientific) at the rate of 0.1 mL/min as image frames were acquired. In single–tethered experiments, the biotinylated ends of DNA were aligned at the barriers while the free ends were extended by flow. In U–shaped DNA experiments, both ends of the DNA molecules were anchored to the lipid, and the molecule adopted a ‘U’ shape under continuous flow when its anchor points were aligned in adjacent wells of the chromium barrier. Additionally, in experiments where nucleosome–bound DNA was used, DNA was first injected into the flow cell in salt–free nucleosome buffer (40 mM Tris–HCl [pH 7.5], 1 mM DTT, 0.5 mg/mL BSA), before the nucleosome buffer was rapidly flushed out and replaced with condensin buffer at the flow rate of 1 mL/min. Dual color imaging of YOYO1–DNA and either ATTO–647N–labeled nucleosomes, or Alexa647–labeled condensin was carried out through additional 640 nm laser excitation (CrystaLaser) and a Dual–View splitter (Optical Insights), allowing simultaneous acquisition in both channels. Photobleaching of CI–Alexa647 was achieved via continuous illumination of the 640 nm laser.

#### Single–molecule data analysis

ND2 files or TIFF image series were imported in ImageJ and saved as TIFF stacks before processing. For single–tethered DNA compaction experiments, a kymograph is generated for each DNA molecule for analysis. Average compaction velocities were estimated by calculating slopes of line segments approximating the trajectory of the DNA free end in the kymograph. Compaction events by definition had positive slopes (velocities). Slippage or disruption of compaction, as well as other events where the DNA free end moved away from the barrier, therefore had negative velocities and were discarded in plotting. Processivity is defined as the total positive displacement between events with negative slopes, or the beginning or end of the trajectory. A compaction event whose observed processivity was considered censored if said event was terminated due to protein dissociation, DNA breakage, free end coming within 1 pixel of the barrier position, or end of data collection period. Survival function of processivity was estimated with the Kaplan–Meier estimator, using the Seaborn library in Python. In U–shaped DNA looping experiments, average velocities of symmetric looping events were measured by estimating the rate at which the looped, overlapping portion of DNA increased in length. Average velocities of asymmetric looping events were measured by manually tracking the displacement of the bright looped DNA signal over time. All distances were initially measured in number of pixels and subsequently converted to μm with the conversion factor of 0.267 μm/pixel, based on the combination of the sensor pixel size and 60X objective. Under the single–molecule experimental conditions specified above, the average extensions of YOYO1–stained naked DNA, nucleosome–bound DNA, and U–shaped DNA were 10.75 ± 0.30 µm (N = 294), 9.11 ± 0.44 µm (N = 112), and 9.54 ± 0.55 µm (N = 91), respectively. Apparent velocities and processivities were converted from physical quantities (µm/s and µm) to more biologically relevant units (bp/s and bp) only to facilitate comparisons with previously reported values. It is important to note that the effective conversion factors (bp/µm) depend not only on the flow rate, but also the DNA geometry, among other factors. Velocity values are reported as median (interquartile range, or IQR) throughout.

#### Histone purification and nucleosome reconstitution

Xenopus H2A (wild–type), H2B (V119C), H3 (C110S), or H4 (wild–type) were expressed in *E. coli* BL21 (DE3) pLysS cells (Novagen, Cat. No. 69451), individually, using a pET3A–based plasmid. Cultures (6 L for each histone) were grown to OD600 ~0.6 at 37°C, and protein expression was induced with 0.5 mM IPTG for 4 hours at 37°C. Cells were harvested at 23°C by centrifugation (3000xg for 10 minutes) and resuspended into 40 mL buffer containing 50 mM Tris (pH 8.0), 10% (w/v) sucrose, 1 mM TCEP, 1 mM PMSF, 1X cOmplete^TM^ Protease Inhibitor Cocktail (Roche, Cat. No. 11697498001) and frozen at –80°C. Thaw the cells on a warm water bath and added 40 ml Tris–washing buffer containing 50 mM Tris [pH 7.5], 100 mM NaCl, 5 mM β–mercaptoethanol, 1mM EDTA, 1X cOmplete^TM^ Protease Inhibitor Cocktail, 1% w/v Triton X–100. Cell suspensions were sonicated for 2 minutes (10s on and 50 s off) at 60% power output. Inclusion bodies were harvested by centrifugation at 20,000xg for 20 minutes at 4°C. The protein pellets were then washed by completely resuspending and spinning 4 times with Tris–washing buffer; Triton X–100 was omitted from the Tris–washing buffer for the final two washes. After the last wash, pellets were stored at –80°C until use. Pellets were mixed with 1 ml dimethyl sulfoxide (DMSO) for 30–60 min at room temperature before adding 40 ml unfolding buffer (20 mM Tris (pH 7.5), 7 M guanidinium–HCL, 10 mM DTT) with stirring for 1 hour at room temperature. Then, suspensions were centrifuged at 20,000xg for 20 minutes to remove any remaining insoluble material. The supernatant was dialyzed against 2L of urea buffer (10 mM Tris (pH 8.0), 7 M urea, 1 mM EDTA, 5 mM β–mercaptoethanol and either 100 mM NaCl for H2A and H2B, or 200 mM NaCl for H3 and H4) in dialysis tubing (3,500 MWCO) overnight at 4°C with two buffer changes. Prior to use, urea buffer was deionized using AG 501–X8 resin (Bio–Rad, Cat. No. 1437424) for 2 hours at room temperature. The dialyzed histones were purified with an Akta FPLC system (GE Healthcare) by loading onto a tandem Q sepharose column (5 ml) followed by a SP sepharose column (5 ml) pre–equilibrated with urea buffer at a flow rate of 0.2 mL/min. The columns were washed with more than 75 mL of urea buffer (100 mM NaCl for H2A and H2B, 200 mM NaCl for H3 and H4), the Q sepharose column was then removed, and the histones bound to the SP sepharose were eluted in urea buffer at 0.2 mL/min using a NaCl gradient to 0.5 M over a total volume of 100 mL. Fractions contain histone were combined and dialyzed against 4 L buffer containing 10mM Tris (pH8.0) and 5 mM β–mercaptoethanol using 3,500 MWCO dialysis tubing with four buffer changes. The initial two rounds were conducted overnight at 4°C, followed by the finial two buffer changes without β–mercaptoethanol with 6–hour intervals. Finally, the histones were frozen in liquid nitrogen and lyophilized in a Labconco FreeZone 1 lyophilizer for 48 hours. The lyophilized histones were stored at –20°C prior to use.

Lyophilized histones (~ 5 mg for each histone) were dissolved into 1 ml unfolding buffer with gentle agitation for 2 hours at room temperature. Histones were mixed at equimolar ratios (use 10–15% more of H2A/H2B relative to H3/H4) and the mixture was diluted to 1 mg/ml with unfolding buffer, and dialyzed against TEB2000 (10 mM Tris [pH 8.0], 1 mM EDTA, 5 mM β–mercaptoethanol, 2 M NaCl) for 48 hours with 4 buffer changes. The dialyzed histones were then concentrated to 1 mL using a 3,000 MWCO spin concentration device (Vivaspin 6, GE Healthcare), and purified on a Superdex S–200 16/60 size exclusion column (GE Healthcare) pre–equilibrated with TEB2000 at a flow rate of 1 mL/min. Fractions contain octamer were combined and concentrated. Equal volume of 100% glycerol was added to the purified octamer solutions (final concentration of 50% glycerol) and stored at –20°C.

Prior to the labeling reaction, potential disulfide bonds in reconstituted histone octamers (H2B V119C) were broken down by incubation with 1 mM Tris(2–carbodxyethyl)phosphine (TCEP; Sigma–Aldrich, Cat. No. C4706) for 20 minutes at room temperature. TECP was then removed from the sample using an Amicon Ultra–0.5 centrifugal filter unit (10,000 MWCO, Millipore Sigma UFC501024) with three ~400–450 μL washes of TE2000 buffer (10 mM Tris [pH 7.0], 1 mM EDTA, 2 M NaCl). 5X molar excess of ATTO–647N maleimide (ATTO–TEC, Cat. No. AD 647N–41) was added to the octamers, and the labeling reaction was incubated at room temperature for 2 hours in the dark. Excess free dye was removed by washing with TE2000 buffer at least 5 times or until flow–through was clear of dye using an Amicon column. Labeling efficiency was determined by both measurement of absorbance (A280) on a NanoDrop 2000 (Thermo Scientific), as well as quantification of SDS–PAGE band intensities scanned on a Typhoon FLA900 (GE Life Sciences) with appropriate settings for fluorescence excitation and emission.

In a 30 μL reconstitution reaction, modified λ–DNA substrate (1.6 nM final concentration) for DNA curtain experiments (BioLDigR for single–tethered and BioLBioR for U–shaped) was mixed with ATTO–647N–labeled histone octamers in TEB2000 buffer with an appropriate DNA:octamer ratio which was adjusted empirically to produce the desired number of nucleosomes per DNA molecule. The reaction was transferred to a home–made dialysis device with 10,000 MWCO SnakeSkin dialysis tubing (Thermo Scientific, Cat. No. 68100) and placed in a beaker containing 100 mL TEB2000 with gentle agitation. Dialysis was carried out overnight at 4 °C with TEB buffer (10 mM Tris [pH 8.0], 1 mM EDTA, pH 8.0, 5 mM β–mercaptoethanol) being continuously added to the beaker at the rate of 0.5 mL/min via a peristaltic pump. After the concentration of NaCl drops below 400 mM, the sample was then dialyzed against 100 mL TEB buffer for another 2 hours. Finally, the reconstitution reaction was retrieved from device and stored at 4°C in the dark.

### Supplemental Movie Legends

**Movie S1. Example of single–tethered DNA compaction by human condensin I.** This video shows a YOYO1–stained single–tethered DNA molecule being fully compacted, from bottom to top, by unlabeled human condensin I (*left*); alongside the kymograph of the compaction (*right*). Compaction was completed at ~300 seconds mark. Vertical and horizontal scale bars represent 2 µm and 60 sec respectively.

**Movie S2. Example of single–tethered DNA compaction by human condensin II.** This video shows a YOYO1–stained single–tethered DNA molecule being fully compacted, from bottom to top, by unlabeled human condensin II (*left*); alongside the kymograph of the compaction (*right*). Compaction was completed at ~415 seconds mark. Vertical and horizontal scale bars represent 2 µm and 60 sec respectively.

**Movie S3. Example of single–tethered DNA compaction by Alexa647–labeled human condensin I.** This video shows a YOYO1–stained single–tethered DNA molecule (green) being compacted, from bottom to top, by Alexa647–labeled human condensin I (magenta); alongside the kymograph of the compaction from the DNA channel, superimposed with tracked trajectory of the labeled CI (*right*). Vertical and horizontal scale bars represent 2 µm and 60 sec respectively.

**Movie S4. Example of single–tethered DNA compaction by Alexa647–labeled human condensin II.** This video shows a YOYO1–stained single–tethered DNA molecule (green) being compacted, from bottom to top, by Alexa647–labeled human condensin II (magenta); alongside the kymograph of the compaction from the DNA channel, superimposed with tracked trajectory of the labeled CII (*right*). Vertical and horizontal scale bars represent 2 µm and 60 sec respectively.

**Movie S5. Symmetric U–shaped DNA compaction by human condensin I.** This video shows a YOYO1–stained U–shaped DNA molecule being compacted symmetrically compacted by unlabeled human condensin I. Scale bar: 1 µm.

**Movie S6. Asymmetric U–shaped DNA compaction by human condensin II.** This video shows a YOYO1–stained single–tethered DNA molecule being asymmetrically compacted by unlabeled human condensin II. Scale bar: 1 µm.

**Movie S7. Nucleosome DNA compaction by human condensin I.** This video shows a YOYO1–stained, nucleosome (magenta)–bound, single–tethered DNA molecule (green) being compacted, from bottom to top, by unlabeled human condensin I; alongside the kymograph of the compaction. Nucleosome–bound DNA was initially fully compacted by ~170 seconds mark, before released and compacted again. Vertical and horizontal scale bars represent 2 µm and 60 sec respectively.

**Movie S8. Nucleosome DNA compaction by human condensin II** This video shows a YOYO1–stained, nucleosome (magenta)–bound, single–tethered DNA molecule (green) being compacted, from bottom to top, by unlabeled human condensin II; alongside the kymograph of the compaction. Nucleosome–bound DNA was initially partially compacted by ~80 seconds mark, before released and fully compacted by ~570 seconds mark. Vertical and horizontal scale bars represent 2 µm and 60 sec respectively.

---

**Fig. S1.**
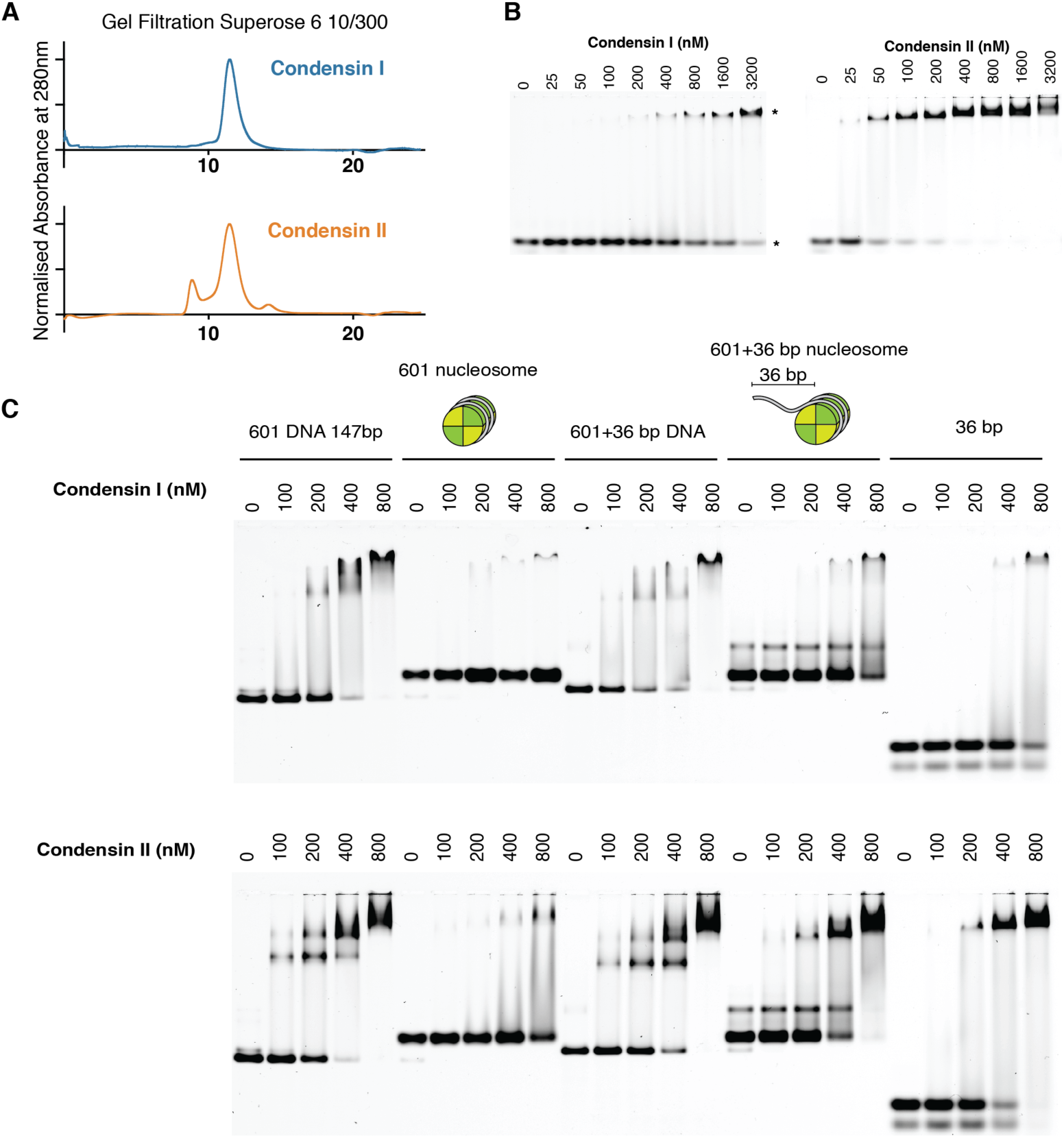
Characterization of human condensin I and II protein and binding to naked and nucleosome–bound DNA. (A) Gel filtration elution profile of condensin I and II using a Superose 6 10/300 column. (B) DNA binding assays for human condensin I (*left*) and II (*right*), using 50 nM of 30 bp dsDNA substrate labeled with Cy5, * indicate free and protein bound DNA. (C) Condensin I (*top*) and II (*bottom*) binding to the 601 nucleosome and nucleosome that has the 601 DNA sequence with an additional 36 bp overhang on one side compared to naked DNA of the same sequence. All DNA labeled with Cy5 and present at 50 nM.

**Fig. S2.**
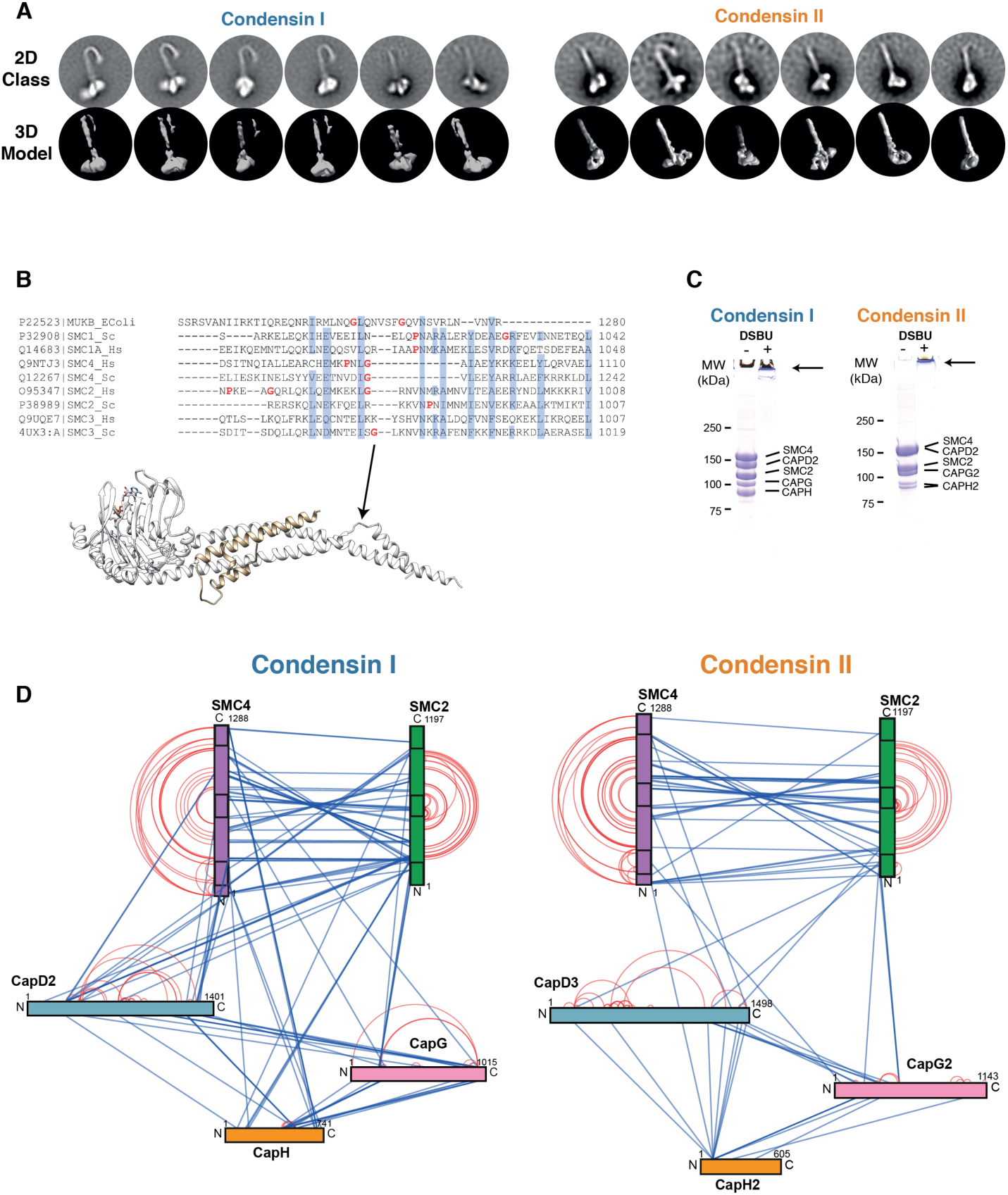
Validation of condensin II model. (A) Comparison between the 2D classes and the 3D model condensin I and II. (B) Sequence alignment of *E. coli*. MukB and *H.sapien* and *S. cerevisae* SMC 1–4 sequence in the region of coiled–coil arm adjacent to ATPase domain. Conserved residues are highlighted blue, with the poor helix forming residues, Glycine and Proline colored red. The structure below is the *S. cerevisae* SMC3–SCC1 crystal structure (PDB ID:4UX3), the arrow indicates the break in helicity due to a proline residue. (C) SDS page gel of condensin I and II samples before and after crosslinking with DSBU, arrow indicates crosslinked sample stuck in well of gel. (D) Network diagram of crosslinks found at least three times in condensin I and II samples. Intra–subunit crosslinks are shown in red, inter–subunit crosslinks are shown in blue. Boxes within SMC2/4 indicate location of ATPase domains at N and C terminus, and central hinge domain.

**Fig. S3.**
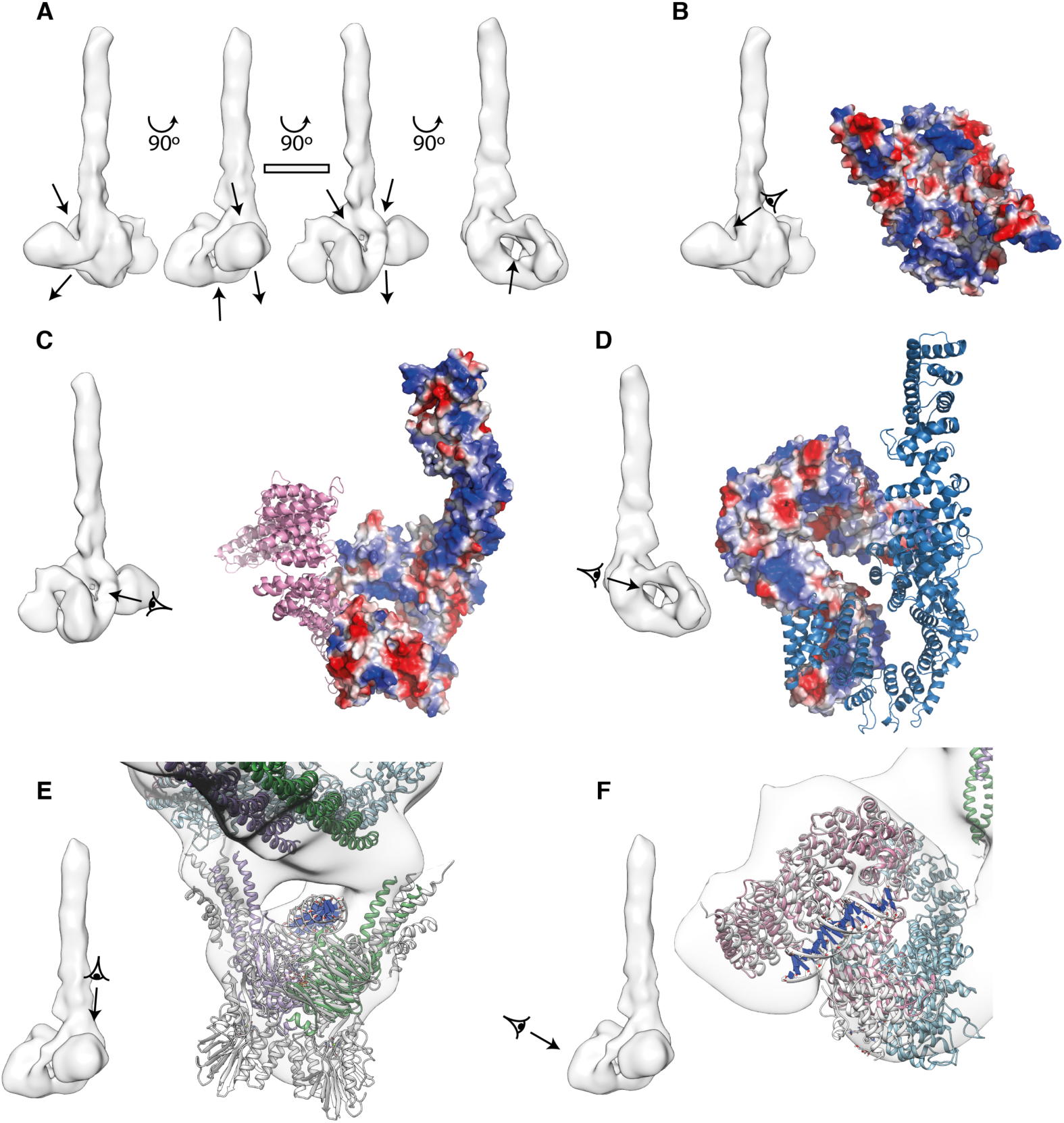
Possible DNA entrapment compartments suggested by condensin II model. (A) 3D model of condensin II, with arrows indicating possible DNA entrapment compartments (scale bar is 10 nm). Electrostatic potential map of interface between SMC2 and 4 ATPase domain (B), of CAP–D3 model, with adjacent CAP–G2 model (C) and of CAP–G2 model, with adjacent CAP–D3 model (D). (E) Overlay of condensin II model with Rad50/MRE11 DNA bound structure (5F3W), which places bound DNA where the hole in density map is located. (F) Overlay of condensin II model with *S. cerevisiae* CAP–G/H DNA structure.

**Fig. S4.**
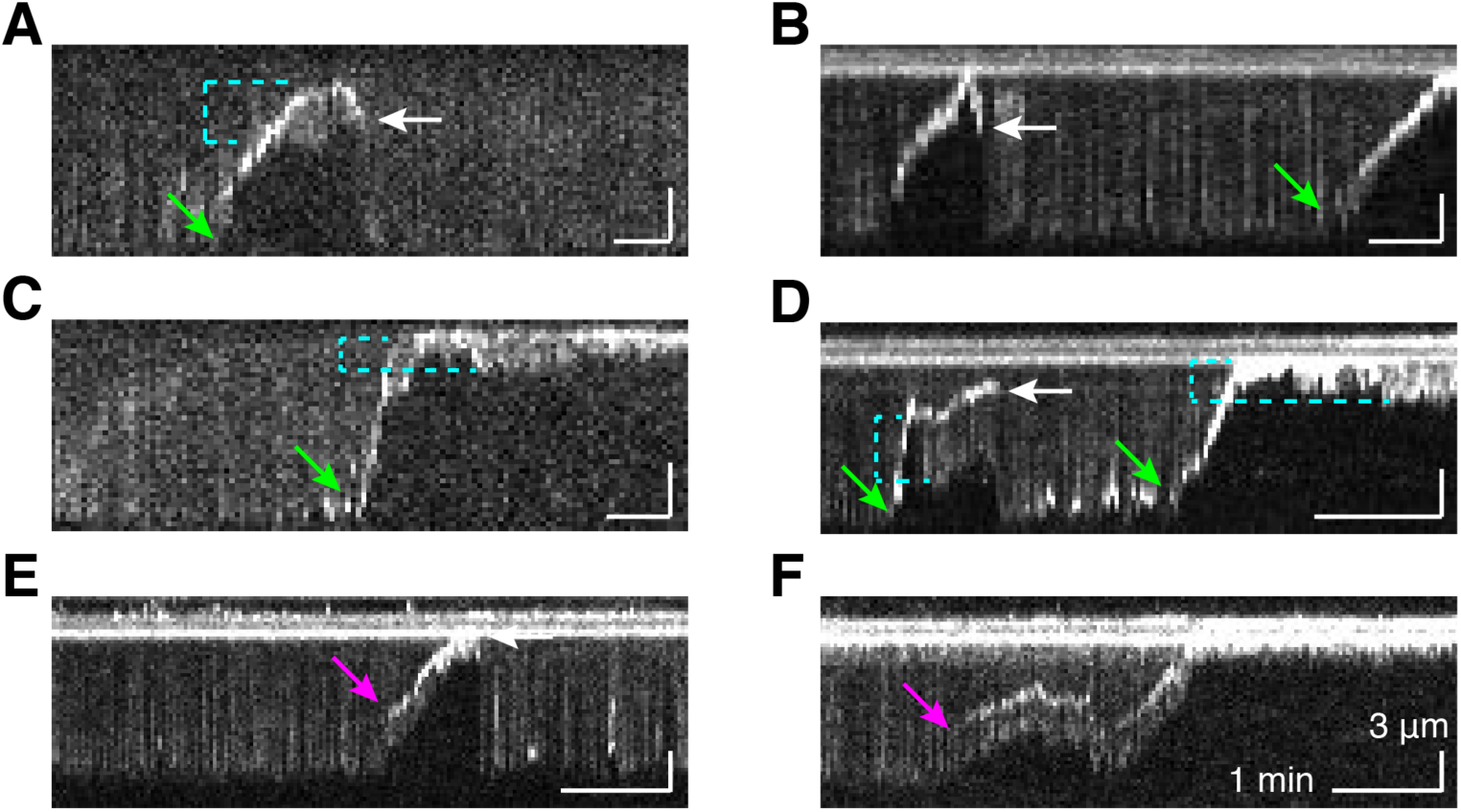
Representative kymographs of single–tethered DNA compaction by human condensin. Green and magenta arrows indicate compaction events that initiated from either the free end or within the internal portion of the DNA molecule, respectively. White arrows indicate sudden and complete release of compacted DNA to its full length. Cyan brackets indicate compacted DNA loops extended by flow.

**Fig. S5.**
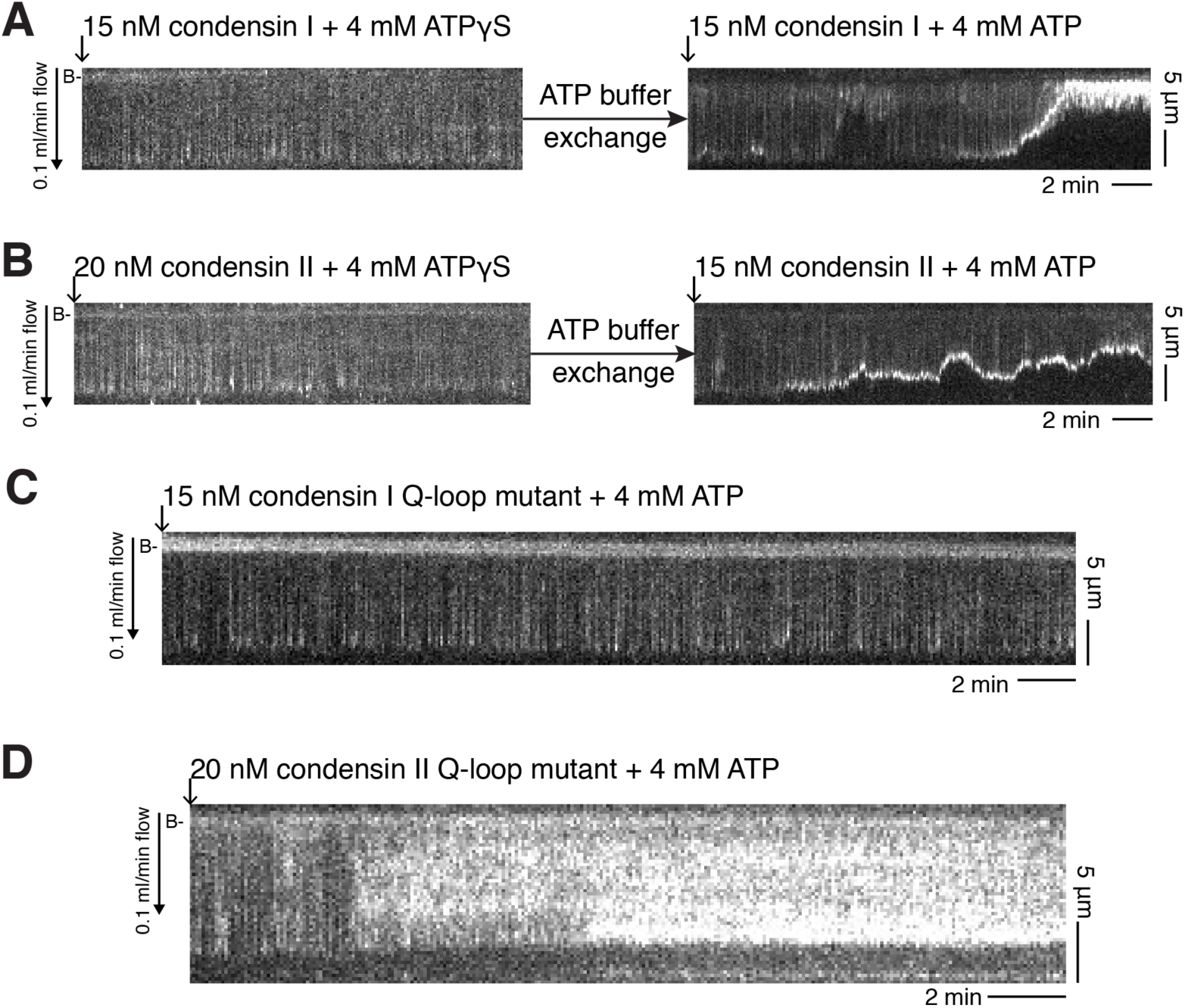
Single–tethered DNA compaction requires ATP hydrolysis. (A) and (B) Representative kymographs of DNA showing no compaction by condensin I or condensin II in the presence of 4 mM ATPγS. After buffer exchange, these same DNA molecules were readily compacted by condensin I and condensin II in the presence of 4 mM ATP. (C) and (D) Representative kymographs of DNA showing no compaction by Q–loop mutants of condensin I or condensin II, in the presence of 4 mM ATP. Note that CII Q–loop mutant was extremely prone to nonspecific interactions with the surface, hence the aberrant appearance of DNA; the brighter signal reflects adherence to the sample chamber surface.

**Fig. S6.**
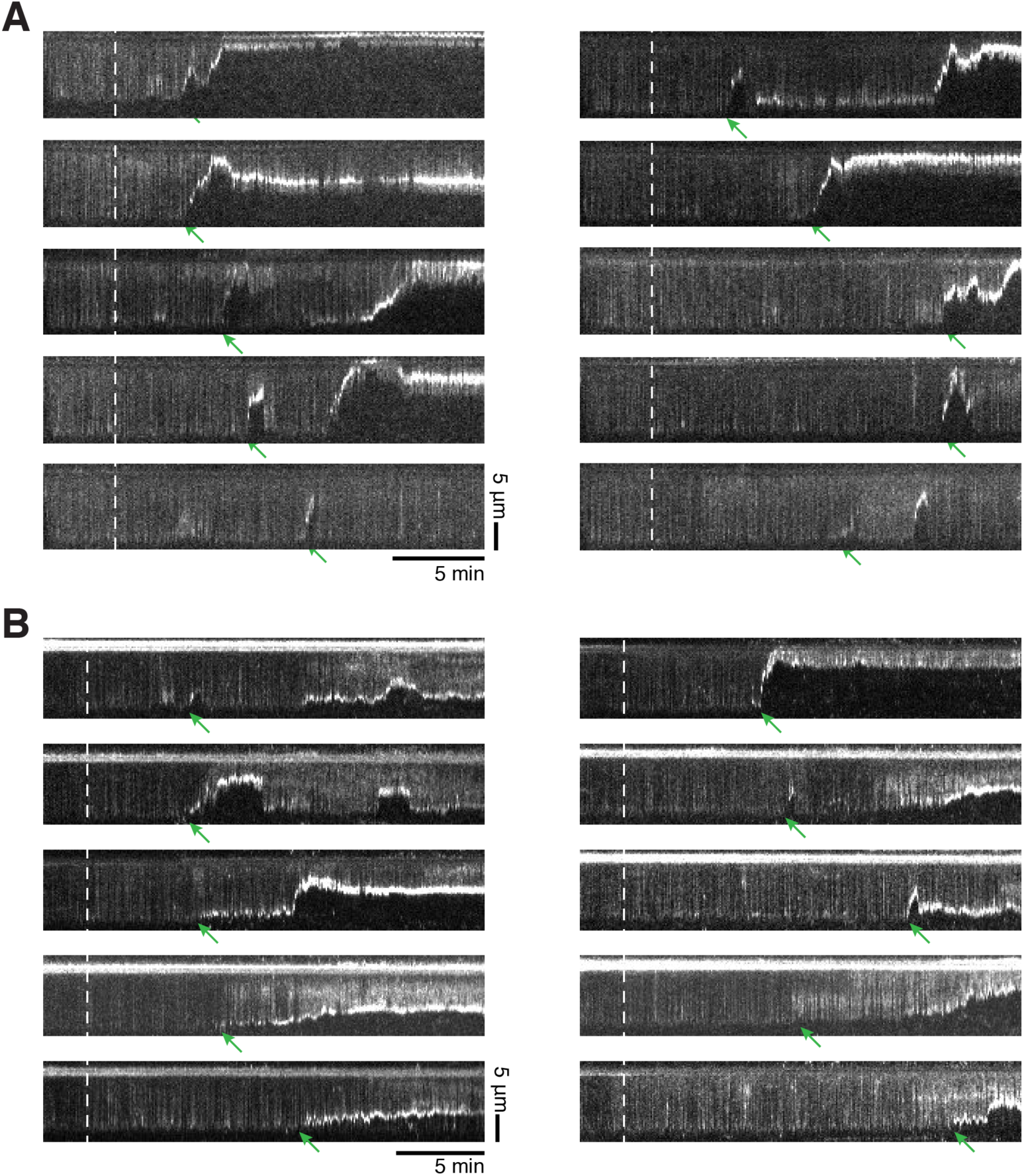
Stochastic initiation of compaction. Representative kymographs of YOYO1–stained single–tethered DNA showing a wide range of initiation start times in compaction events by condensin I (A) and II (B). White dashed lines mark the proteins’ predicted times of arrival in flow cells. Green arrows indicate initiations of the first compaction events on each DNA molecule.

**Fig. S7.**
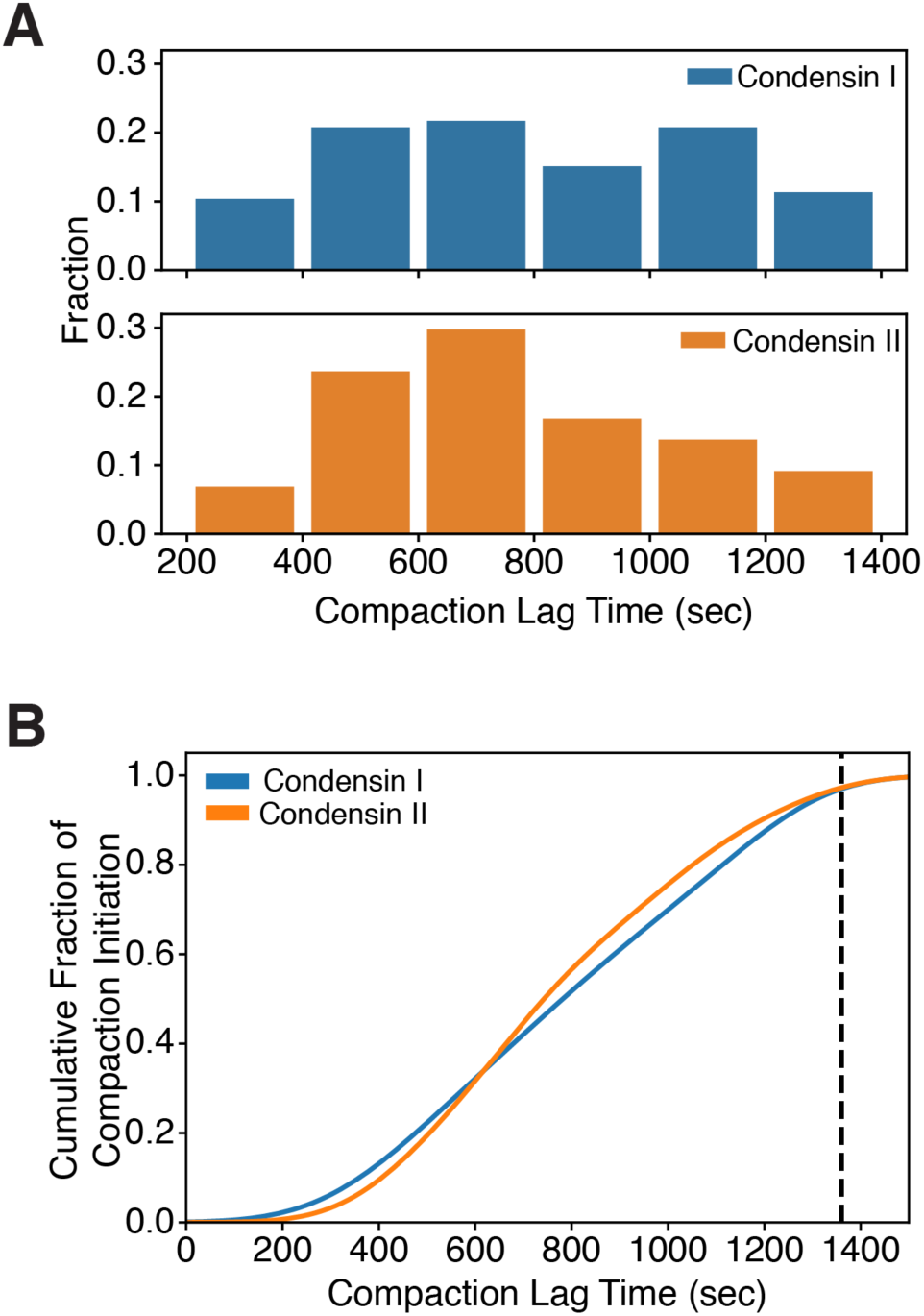
Lag times in initiation of compaction by condensin I and II. (A) Distribution of compaction lag times for the first compaction events on individual DNA molecules. (B) Cumulative fraction of initiation of DNA compaction events.

**Fig. S8.**
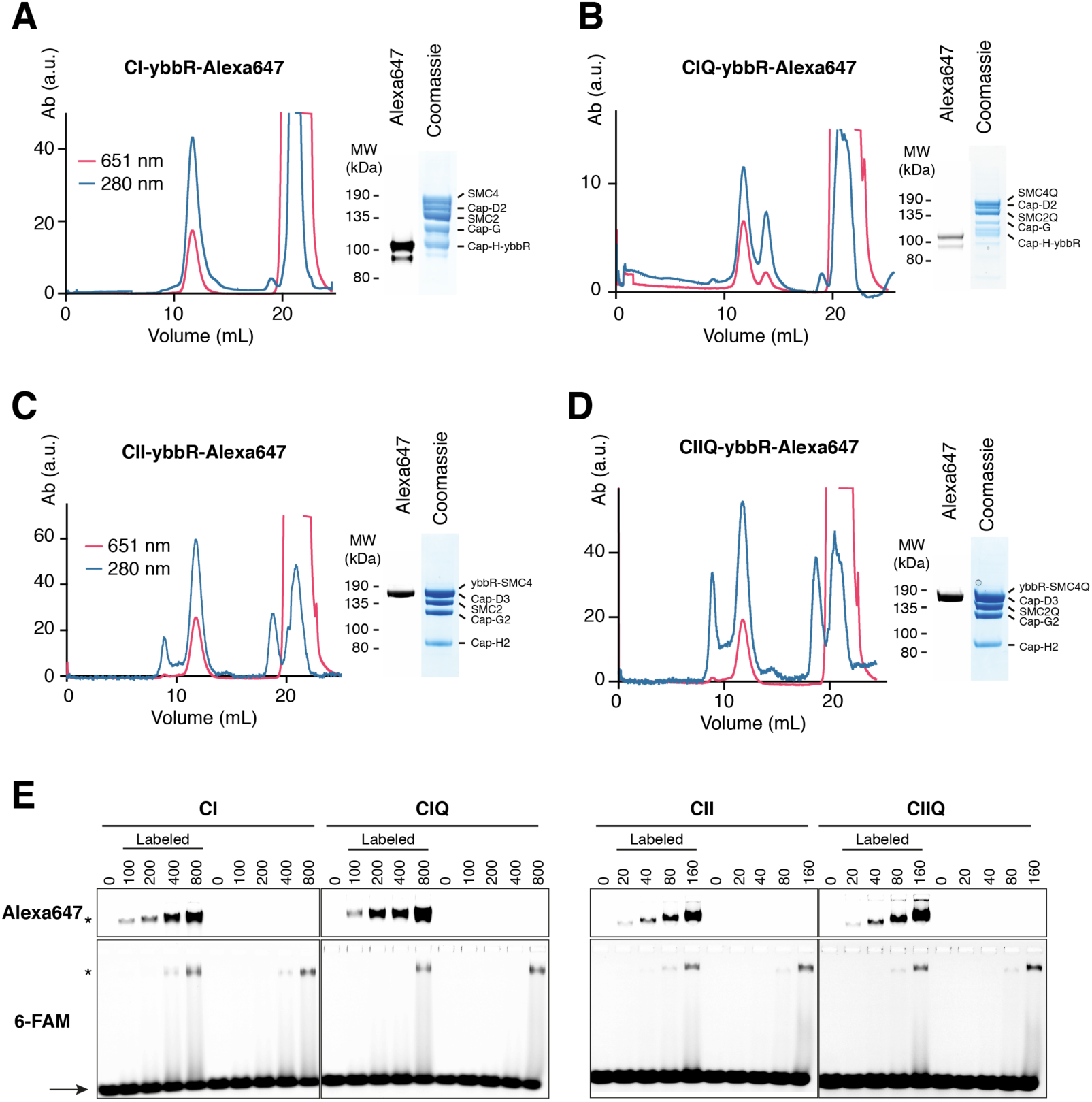
Purification of Alexa–647–labeled condensin I and II. (A) Chromatogram of gel filtration with Superose 6 Increase 10/300 column of labeled CI, showing absorption at the excitation wavelength of Alexa–647 (651 nm) and protein absorption (280 nm). Protein peak elutes at ~11 mL, while excess SFP and unconjugated fluorophore elutes after 20 mL. Insert shows SDS page sample from protein peak, imaged for Alexa–647, followed by Coomassie staining. (B, C and D) As in (A) for CI Q loop mutant, CII and CII Q loop mutant respectively. (E) EMSA assay using 6–FAM labeled 30bp DNA, and Alexa–647 labeled and unlabeled CI, CIQ, CII and CIIQ from left to right respectively. Arrow indicates free DNA, * indicated shifted DNA.

**Fig. S9.**
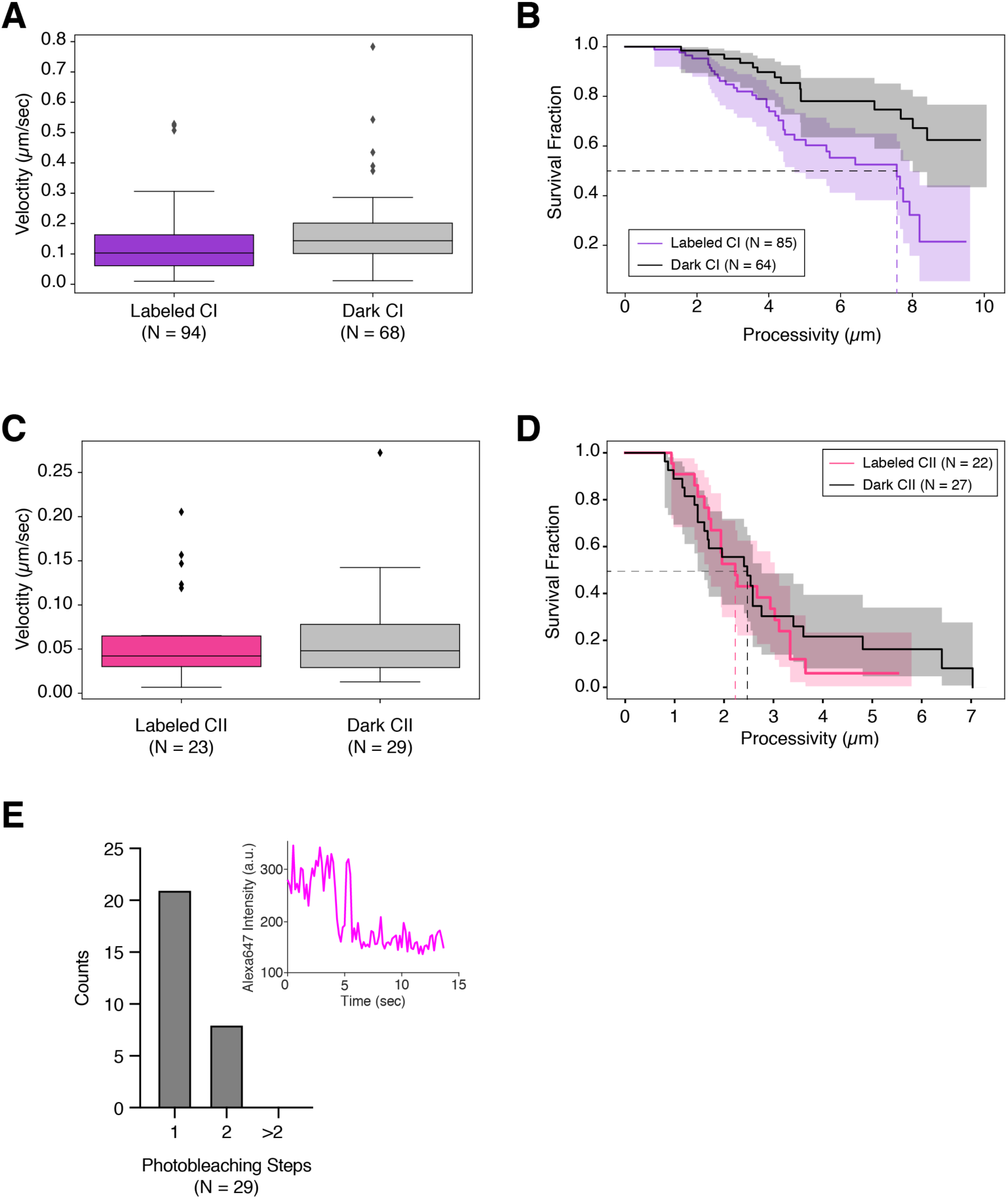
Compaction of single–tethered naked DNA by Alexa–647–labeled condensin I and II. (A) and (B) Compaction velocities (labeled CI median (IQR): 0.103 (0.102) µm/s or ~465 bp/s; dark CI median (IQR): 0.143 (0.100) µm/s, or ~645 bp/s) and processivities (labeled CI half–life: 7.56 µm, or ~34.1 bp; dark CI half–life: n.d.), respectively, of YOYO–1 signal puncta co–localized and tracking with labeled or dark CI–Alexa647. (C) and (D) Compaction velocities (labeled CII median (IQR): 0.042 (0.035) µm/s, or ~189 bp/s; dark CII median (IQR): 0.048 (0.049) µm/s, or ~217 bp/s) and processivities (labeled CII half–life: 2.22 µm, or ~10.0 kbp; dark CII half–life: 2.47 µm, or ~11.1 kbp), respectively, of YOYO–1 signal puncta co–localized and tracking with labeled or dark CII–Alexa647. (E) Histogram of the number of photobleaching steps for CI–Alexa647 complexes that were involved in DNA compaction. Inset: Representative one–step photobleaching trace of CI–Alexa647.

**Fig. S10.**
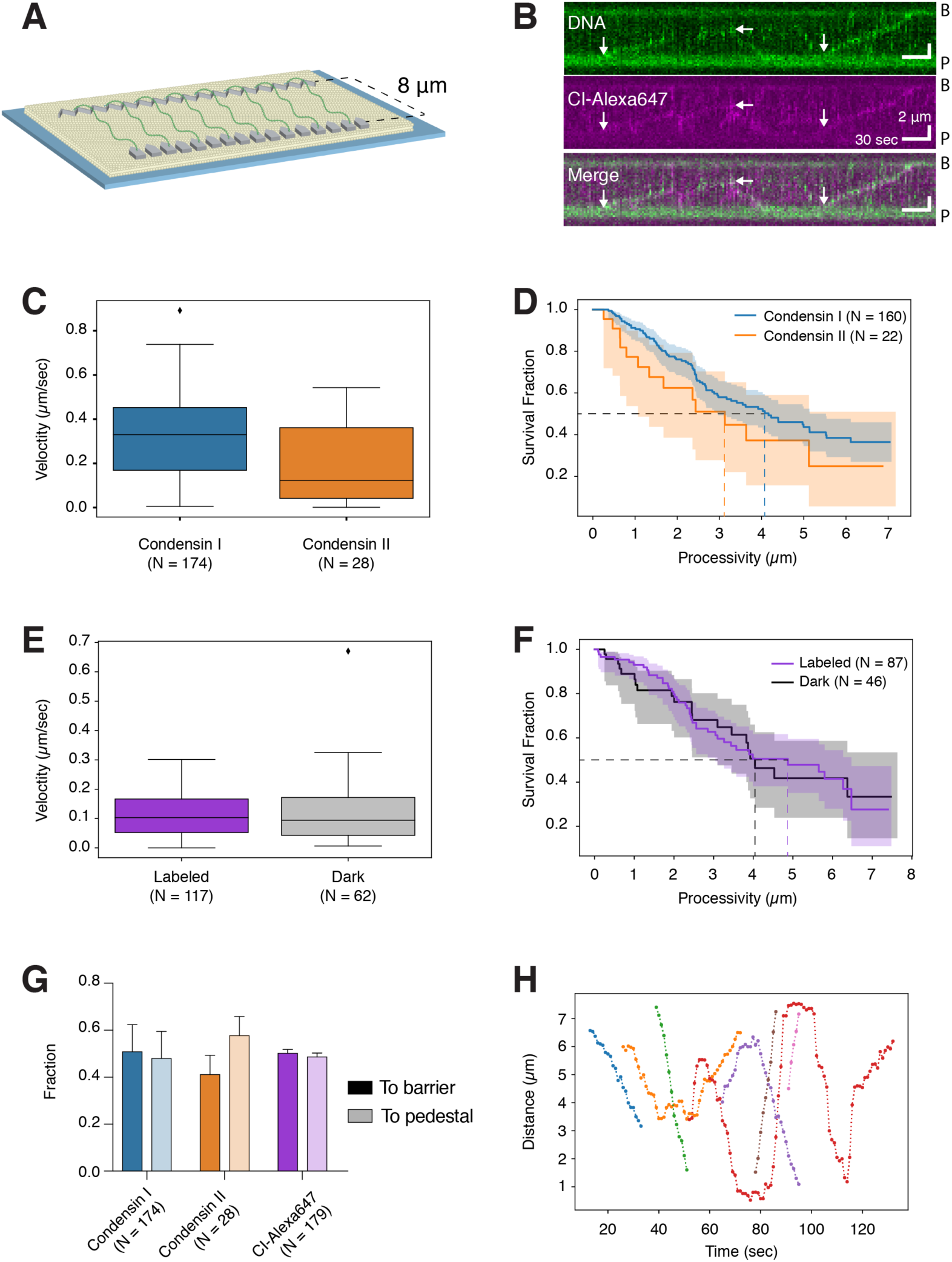
Translocation of unlabeled and labeled CI and CII on 8 µm double–tethered λ–DNA. (A) Schematic of double–tethered DNA curtain assay. (B) Representative kymographs of Alexa647–labeled human CI translocating on 8 µm double–tethered λ–DNA. White arrows indicate starting points of individual translocation events. (C) and (D) Translocation velocities (CI median (IQR): 0.33 (0.28) µm/s, or ~2002 bp/s; CII median (IQR): 0.12 (0.32) µm/s, or ~746 bp/s) and processivities (CI half–life: 4.16 µm, or ~25.2 kbp; CII half–life: 3.13 µm, or ~19.0 kbp), respectively, of YOYO–1 signal puncta along double–tethered DNA molecules in the presence of unlabeled CI or CII. (E) and (F) Translocation velocities (labeled CI median (IQR): 0.103 (0.114) µm/s, or ~624 bp/s, dark CI median: 0.094 (0.129) µm/s, or ~570 bp/s) and processivities (labeled CI half–life: 4.87 µm, or ~29.5 kbp; dark CI half–life: 4.04 µm, or ~24.5 kbp), respectively, of YOYO–1 signal puncta along double–tethered DNA molecules co–localized and tracking with labeled or dark CI–Alexa647. (G) Fractions of unlabeled or labeled condensin translocation events towards the barrier (dark colors) or the pedestal (light colors). (H) Representative single particle tracking trajectories of unlabeled CsI translocation events.

**Fig. S11.**
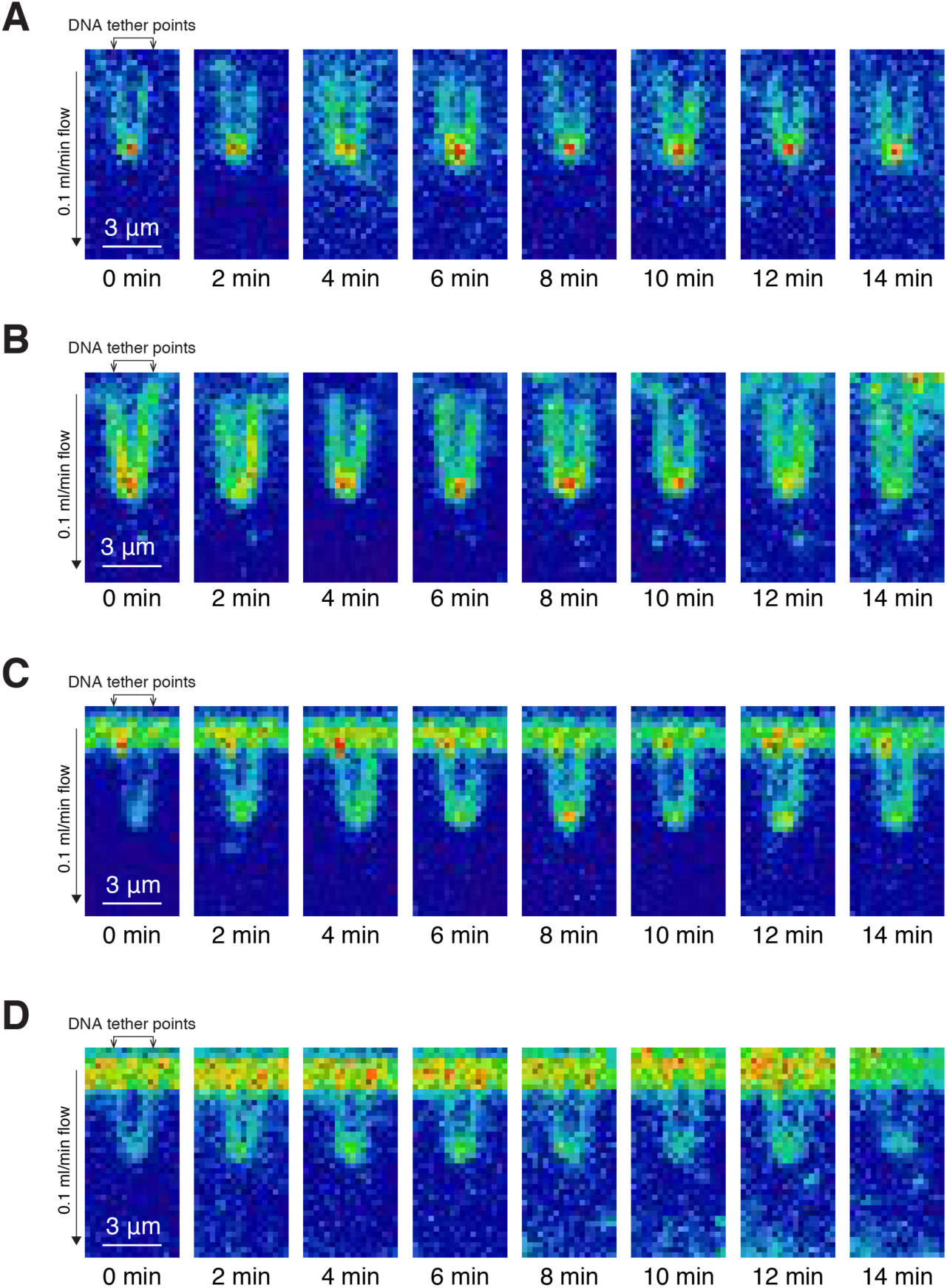
Looping and compaction of U–shaped DNA requires ATP hydrolysis. (A) and (B) Representative snapshots of U–shaped DNA showing no looping or compaction by condensin I or condensin II in the presence of 4 mM ATPγS. (C) and (D) Representative snapshots of U–shaped DNA showing no looping or compaction by Q–loop mutants of condensin I or condensin II, in the presence of 4 mM ATP.

**Fig. S12.**
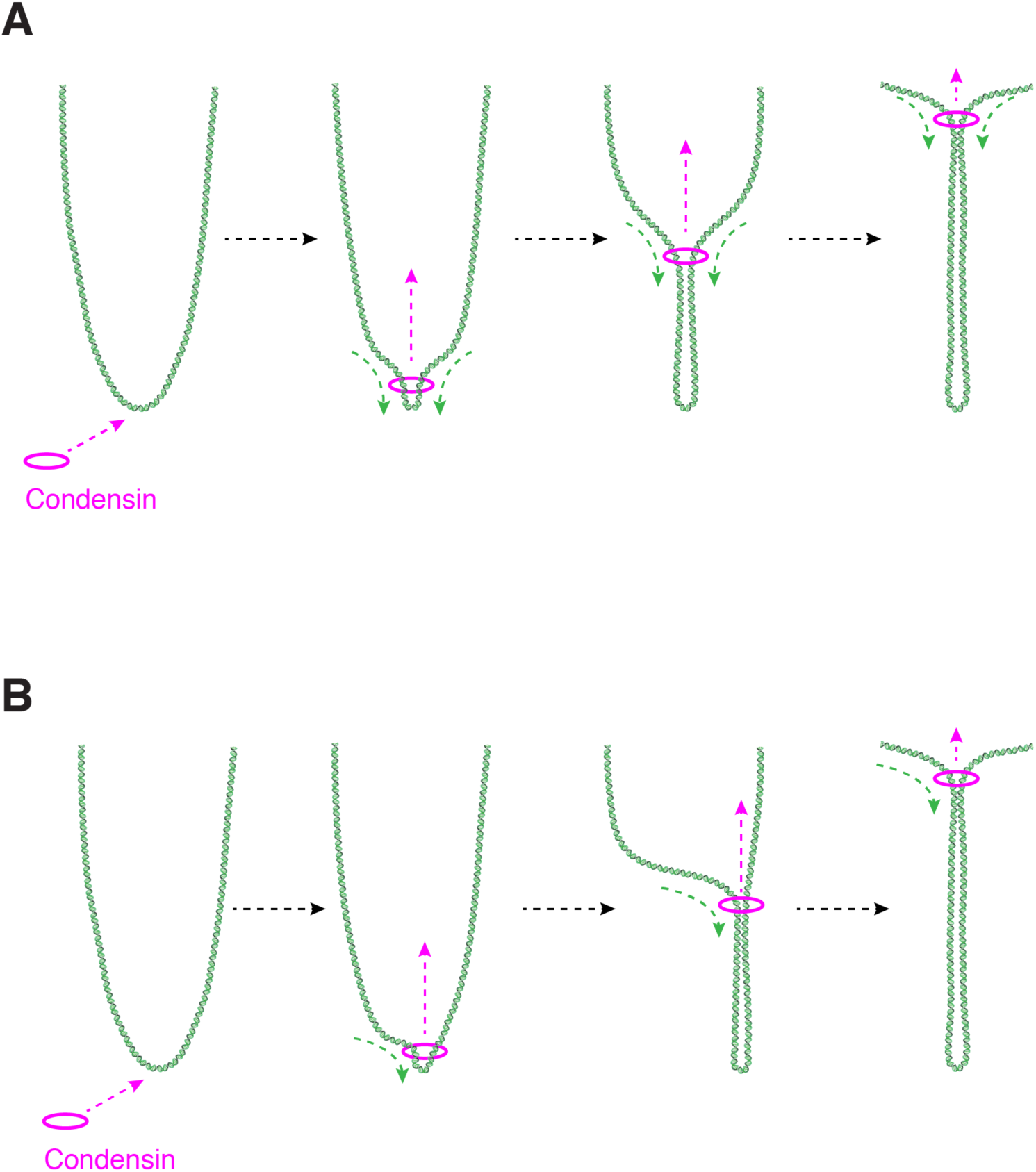
Schematics of symmetric and asymmetric looping of U–shaped DNA by human condensins. (A) Schematics of symmetric looping of U–shaped DNA by human condensins. (B) Schematics of asymmetric looping of U–shaped DNA by human condensins.

**Fig. S13.**
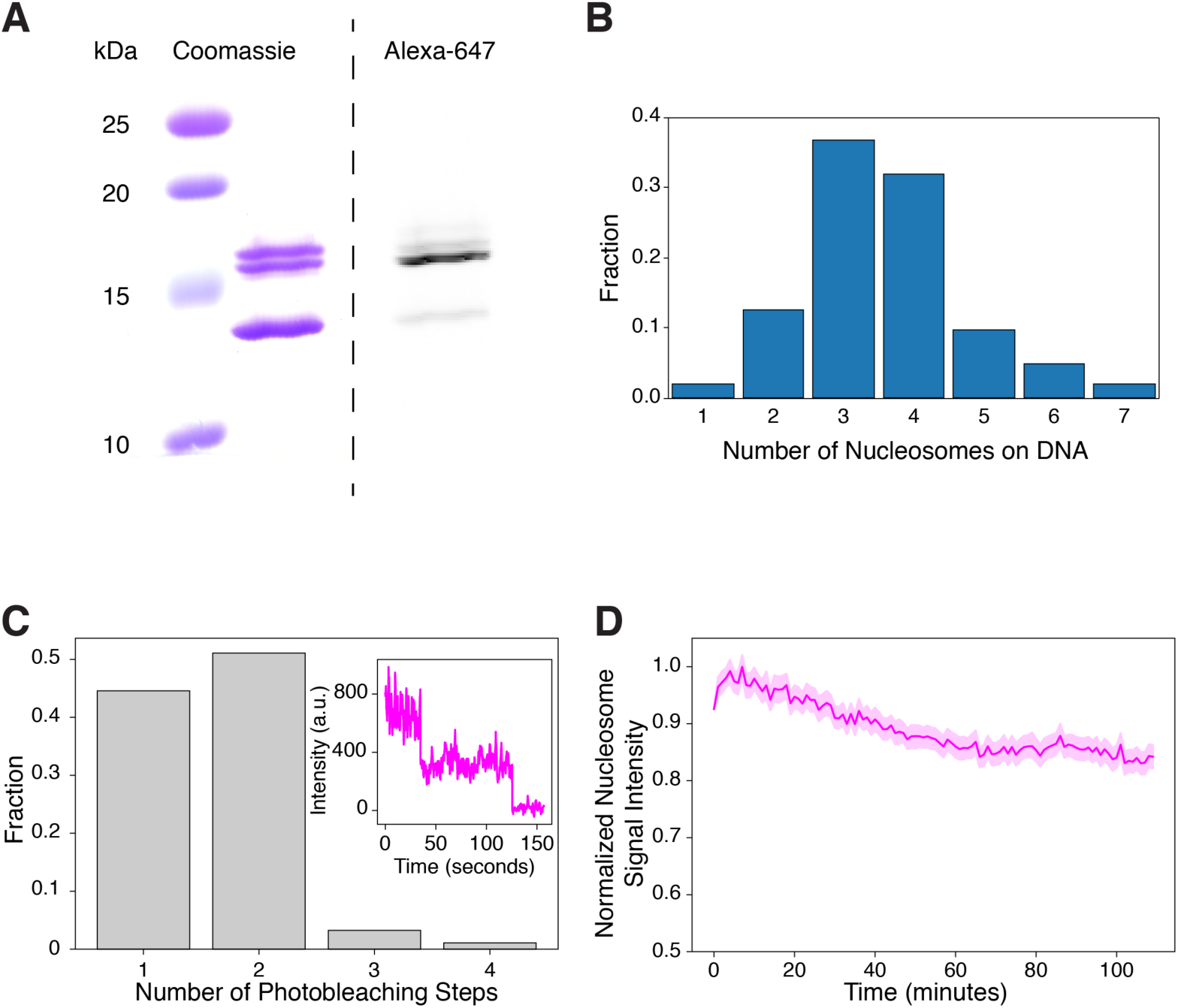
Xenopus nucleosome reconstitution controls. (A) Coomassie staining and ATTO–647N scan of SDS–PAGE of ATTO–647N–labeled xenopus histone octamers. (B) Distribution of numbers of reconstituted nucleosomes on DNA molecules (N=103). (C) Distribution of numbers of photobleaching steps at ATTO–647N signal puncta on DNA (N=92). Inset: representative trace showing two–step photobleaching. (D) Normalized nucleosome signal intensity over time, under single–molecule DNA curtain assay conditions. Shaded region indicates standard deviation across DNA molecules (N=190).

**Fig. S14.**
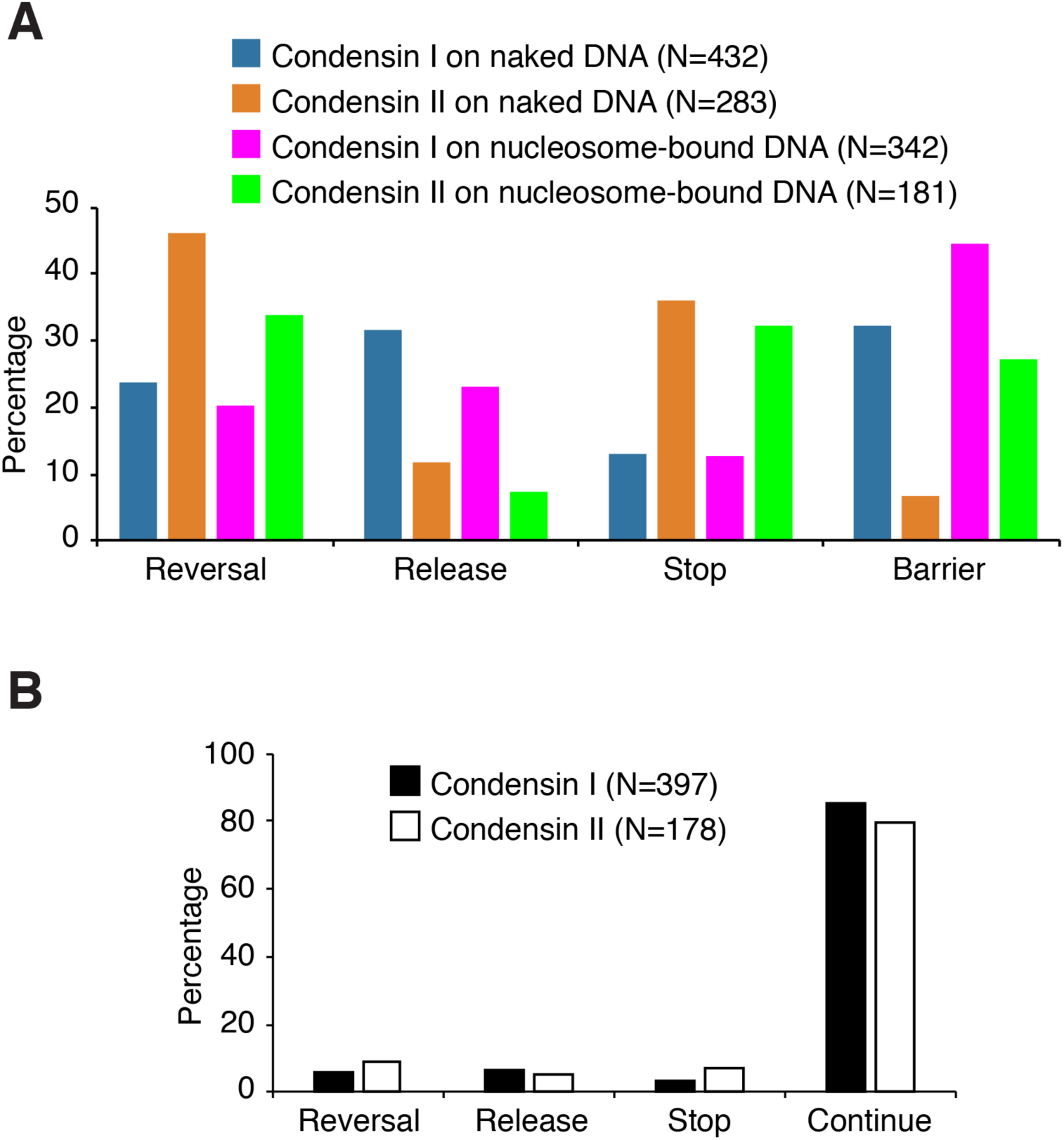
Behavior of human condensin in single–tethered DNA compaction events. (A) Percentages of event types that resulted in termination of DNA compaction events by condensin on either naked DNA or nucleosome–bound DNA. (B) Percentages of outcomes in collisions between condensins and individual nucleosomes during DNA compaction events.

**Fig. S15.**
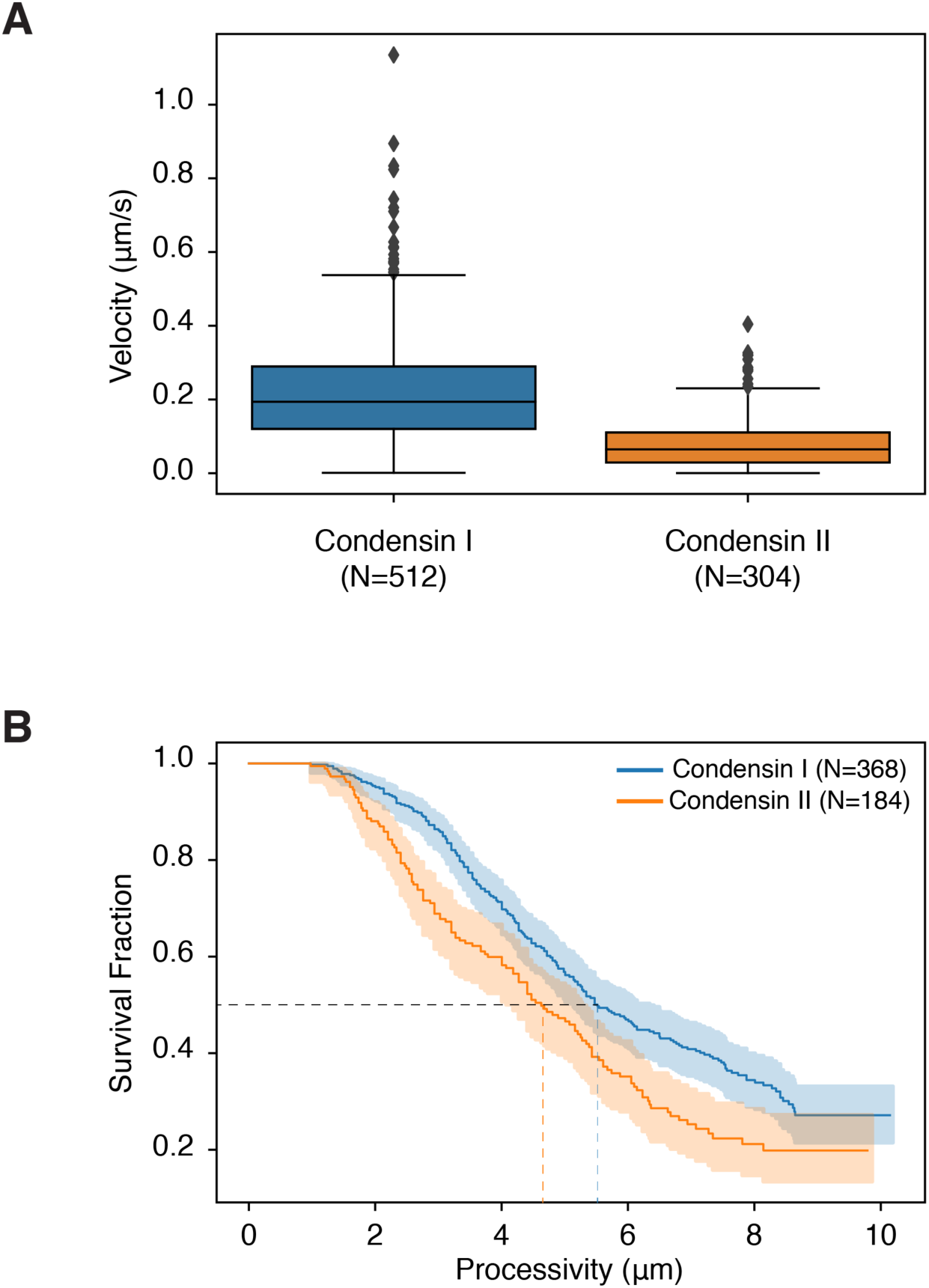
Compaction velocities and processivities of human condensins on single–tethered nucleosome–bound DNA. (A) Box plot of compaction velocities of condensin I and II on nucleosome–bound DNA. Median velocities (IQR) for CI and CII on nucleosome–bound DNA are 0.194 (0.169) µm/s, or ~1032 bp/s; and 0.065 (0.082) µm/s, or ~345 bp/s, respectively. (B) Kaplan–Meier estimated survival functions of compaction processivities on nucleosome–bound DNA. Shaded areas indicate 95% confidence intervals. Median processivities for CI and CII on nucleosome–bound DNA are 5.52 µm (or ~29.4 kbp) and 4.63 µm (or ~24.6 kbp), respectively.

**Table S1.**
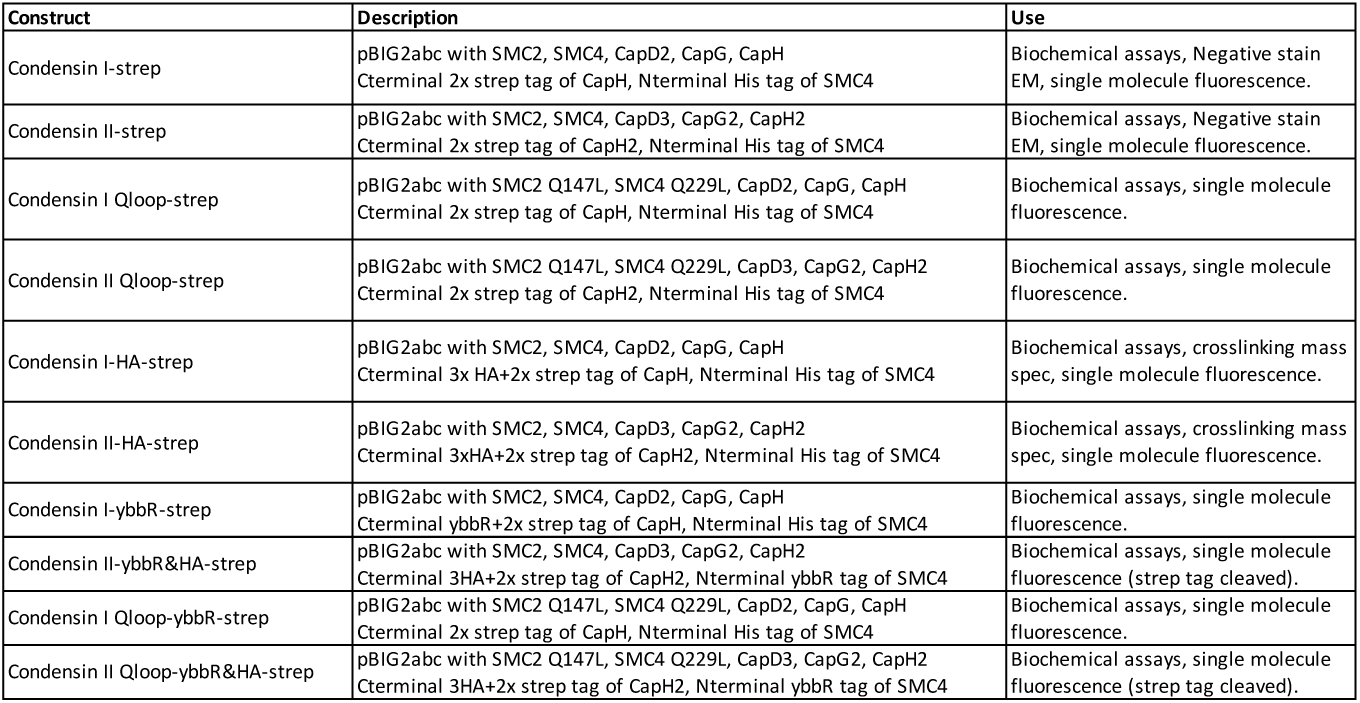
Human condensin constructs purified.

| Construct | Description | Use |
| --- | --- | --- |
| Condensin I-strep | pBIG2abc with SMC2, SMC4, CapD2, CapG, CapH<br>Cterminal 2x strep tag of CapH, Nterminal His tag of SMC4 | Biochemical assays, Negative stain EM, single molecule fluorescence. |
| Condensin II-strep | pBIG2abc with SMC2, SMC4, CapD3, CapG2, CapH2<br>Cterminal 2x strep tag of CapH2, Nterminal His tag of SMC4 | Biochemical assays, Negative stain EM, single molecule fluorescence. |
| Condensin I Qloop-strep | pBIG2abc with SMC2 Q147L, SMC4 Q229L, CapD2, CapG, CapH<br>Cterminal 2x strep tag of CapH, Nterminal His tag of SMC4 | Biochemical assays, single molecule fluorescence. |
| Condensin II Qloop-strep | pBIG2abc with SMC2 Q147L, SMC4 Q229L, CapD3, CapG2, CapH2<br>Cterminal 2x strep tag of CapH2, Nterminal His tag of SMC4 | Biochemical assays, single molecule fluorescence. |
| Condensin I-HA-strep | pBIG2abc with SMC2, SMC4, CapD2, CapG, CapH<br>Cterminal 3x HA+2x strep tag of CapH, Nterminal His tag of SMC4 | Biochemical assays, crosslinking mass spec, single molecule fluorescence. |
| Condensin II-HA-strep | pBIG2abc with SMC2, SMC4, CapD3, CapG2, CapH2<br>Cterminal 3xHA+2x strep tag of CapH2, Nterminal His tag of SMC4 | Biochemical assays, crosslinking mass spec, single molecule fluorescence. |
| Condensin I-ybbR-strep | pBIG2abc with SMC2, SMC4, CapD2, CapG, CapH<br>Cterminal ybbR+2x strep tag of CapH, Nterminal His tag of SMC4 | Biochemical assays, single molecule fluorescence. |
| Condensin II-ybbR&HA-strep | pBIG2abc with SMC2, SMC4, CapD3, CapG2, CapH2<br>Cterminal 3HA+2x strep tag of CapH2, Nterminal ybbR tag of SMC4 | Biochemical assays, single molecule fluorescence (strep tag cleaved). |
| Condensin I Qloop-ybbR-strep | pBIG2abc with SMC2 Q147L, SMC4 Q229L, CapD2, CapG, CapH<br>Cterminal 2x strep tag of CapH, Nterminal His tag of SMC4 | Biochemical assays, single molecule fluorescence. |
| Condensin II Qloop-ybbR&HA-strep | pBIG2abc with SMC2 Q147L, SMC4 Q229L, CapD3, CapG2, CapH2<br>Cterminal 3HA+2x strep tag of CapH2, Nterminal ybbR tag of SMC4 | Biochemical assays, single molecule fluorescence (strep tag cleaved). |

**Table S2.**
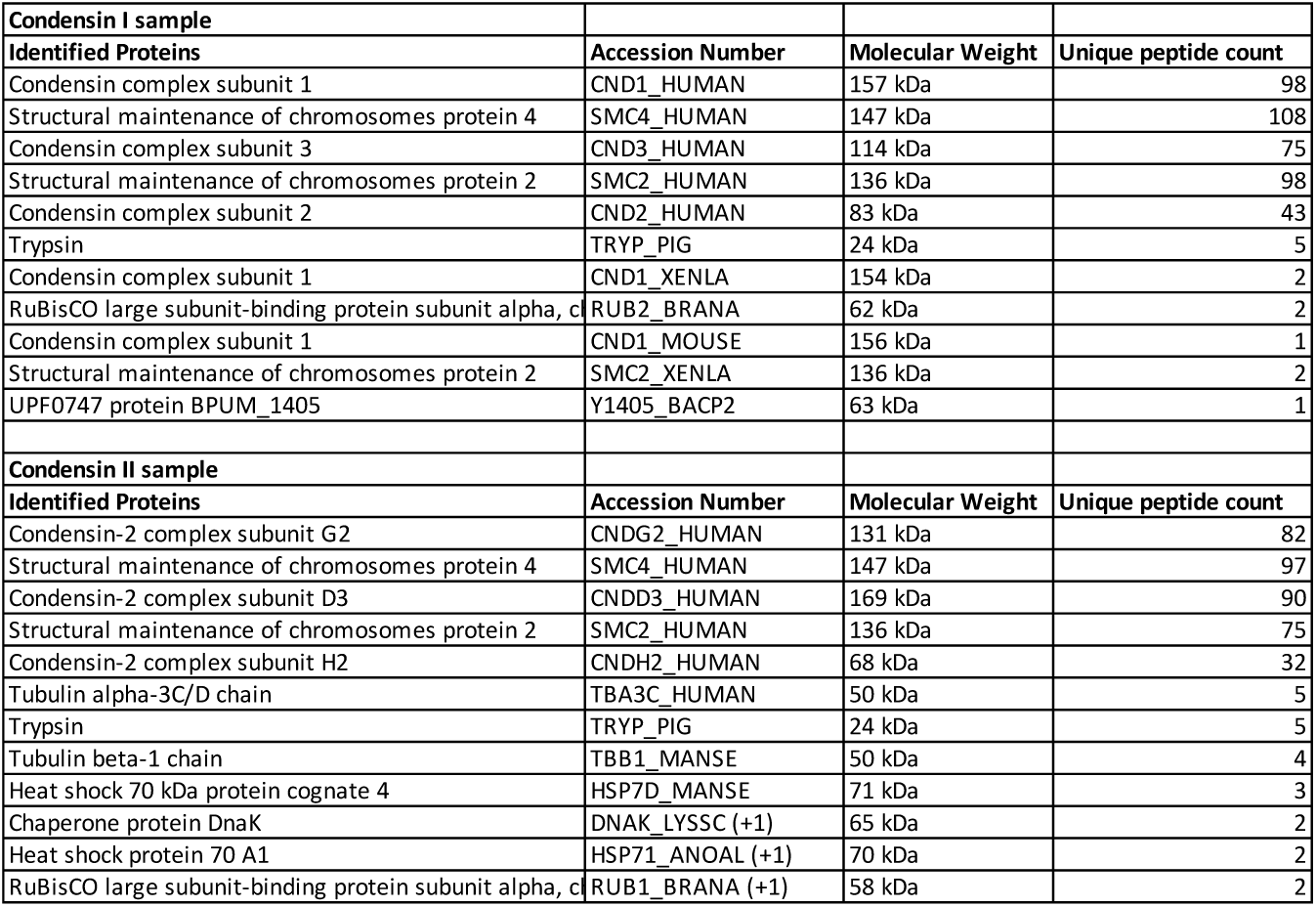
MS/MS protein identification of purified condensin I and II.

**Table S3.**
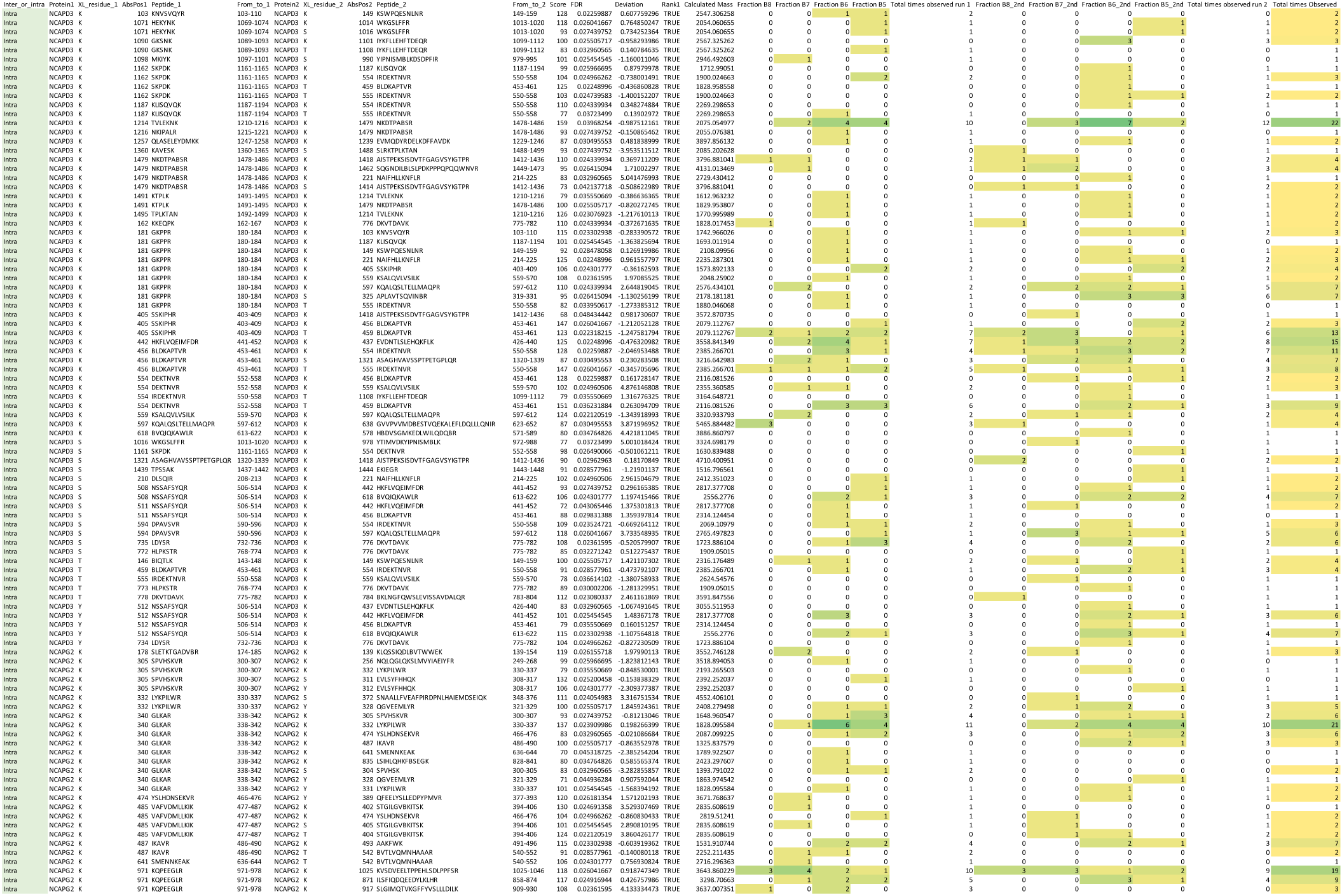
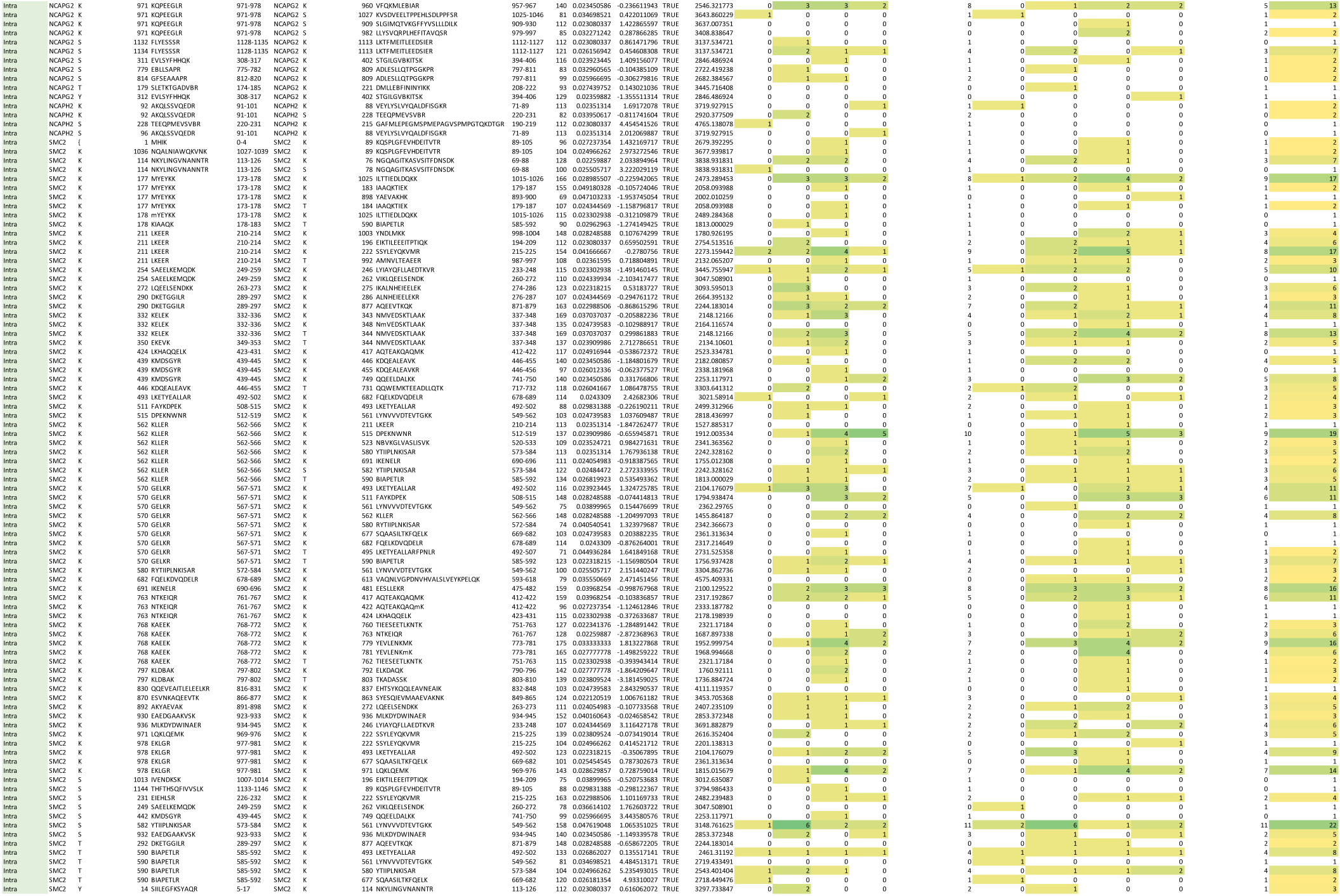
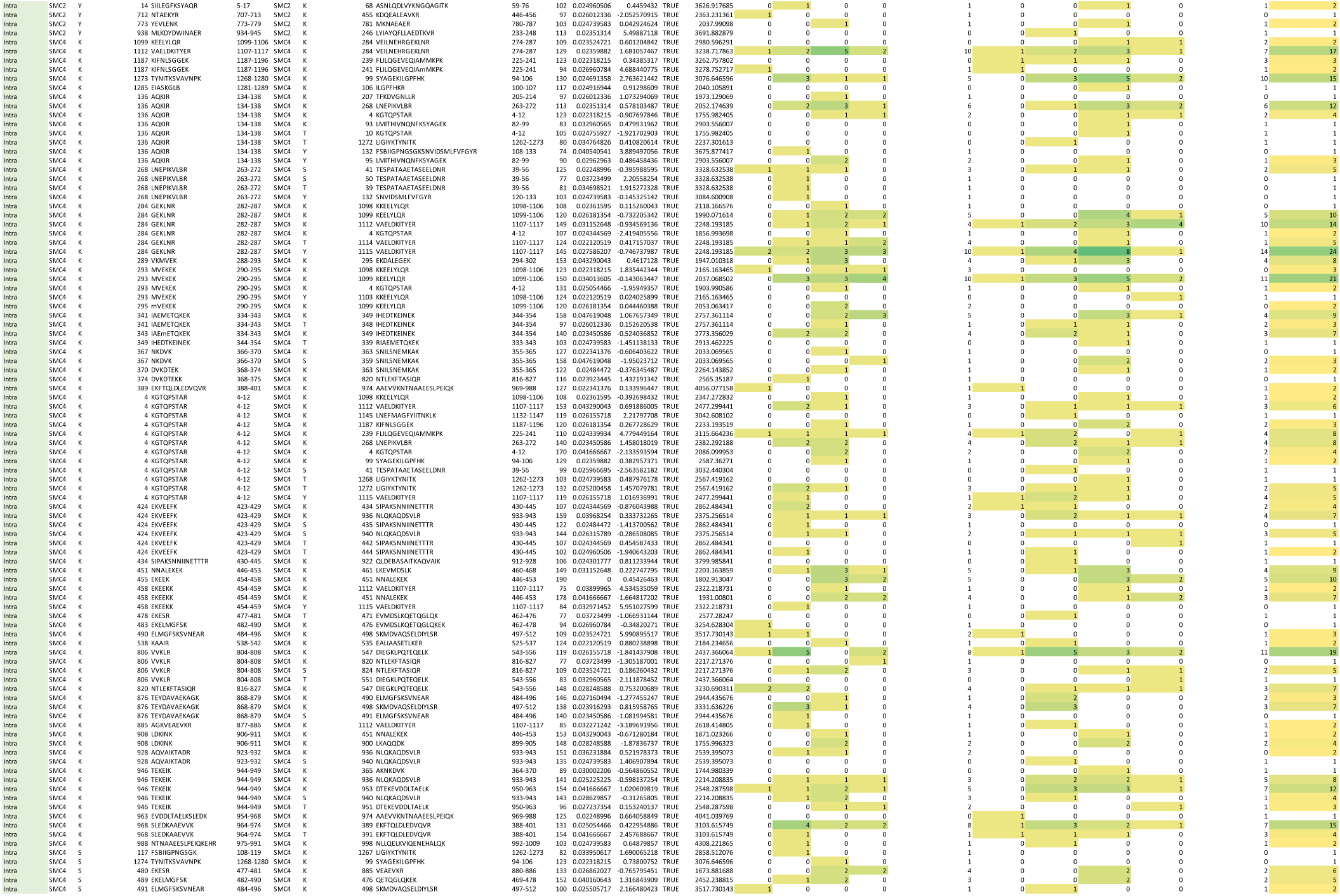
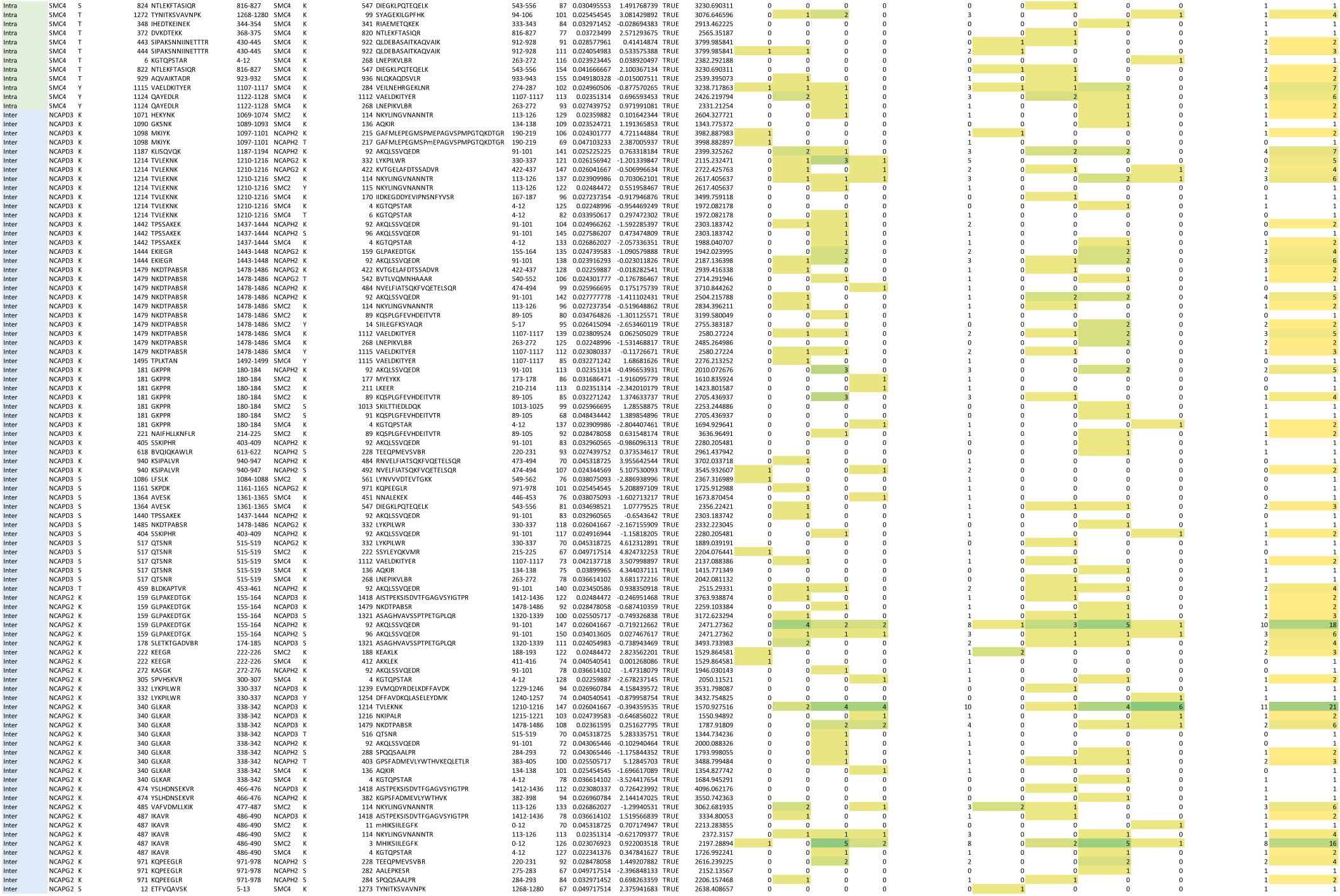
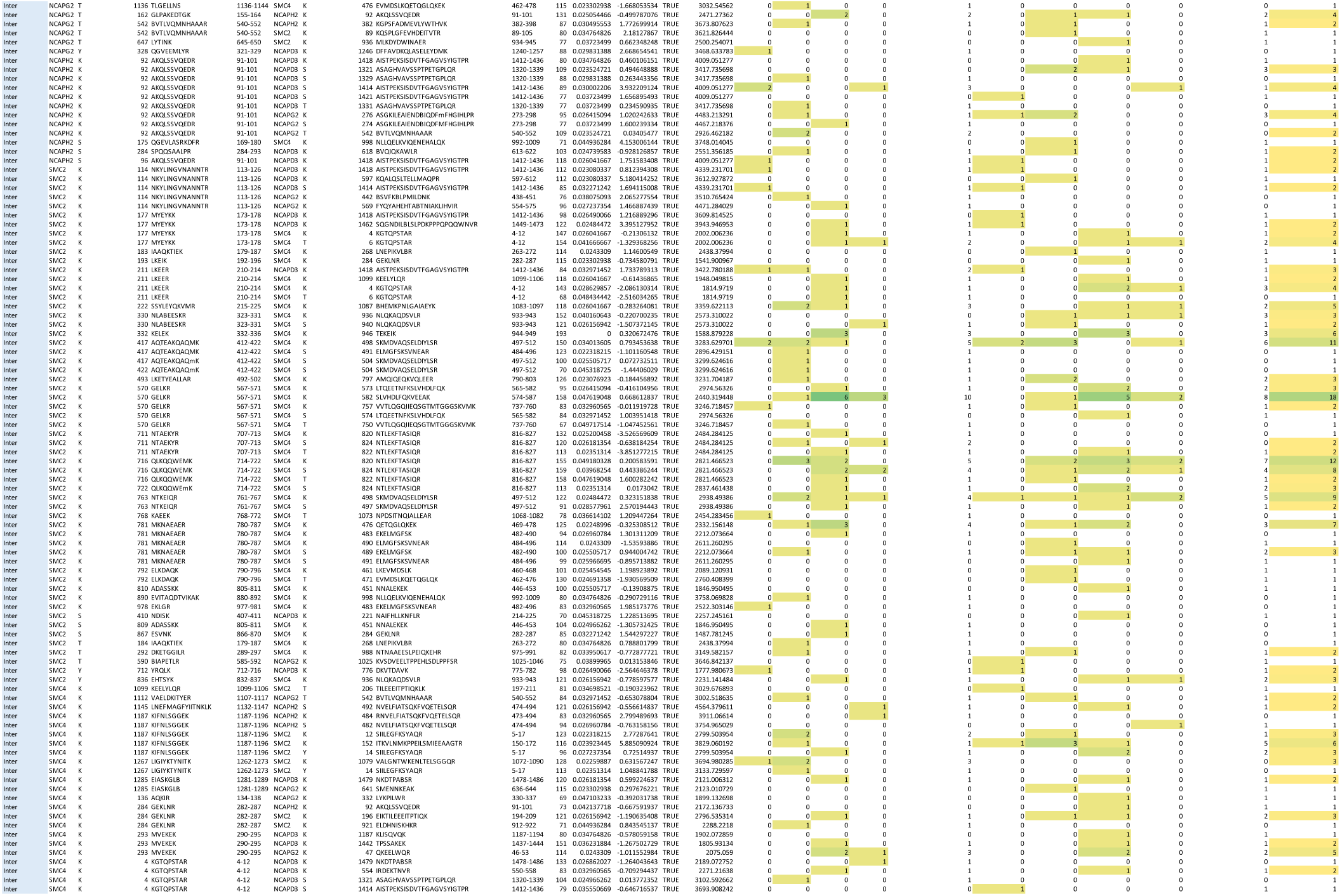
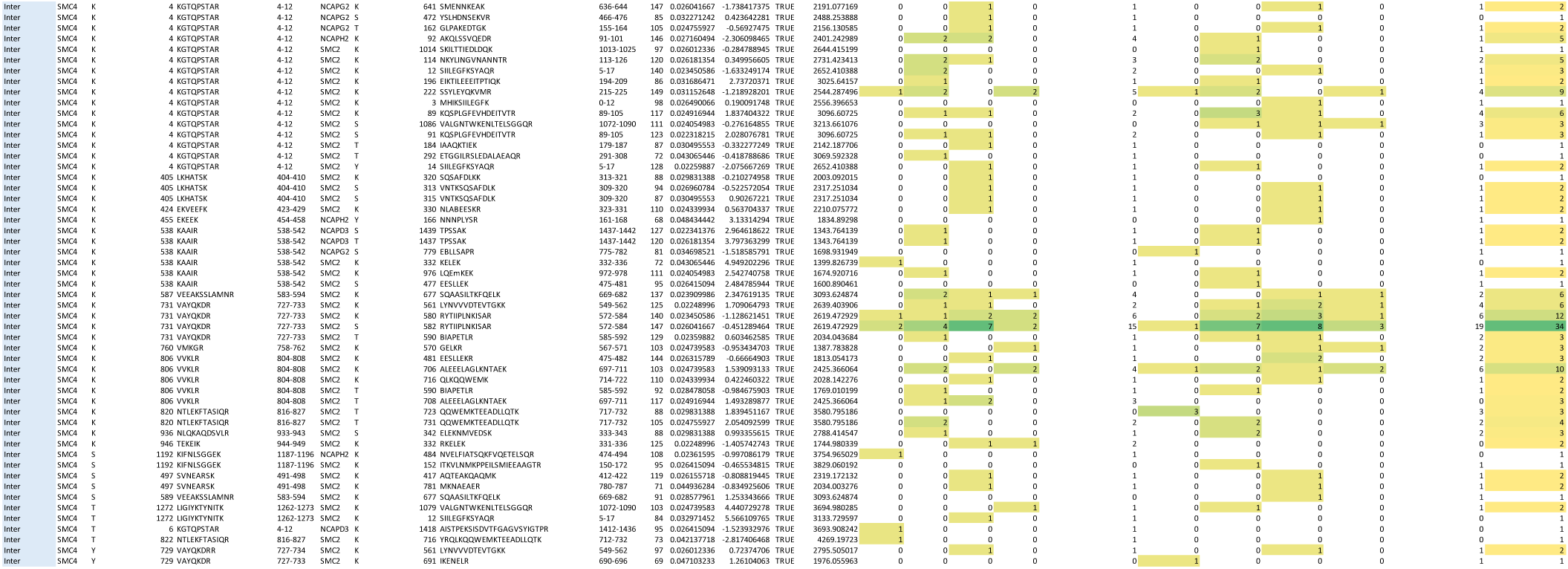
All crosslinks identified by mass spectroscopy in condensin I and II.

